# Role of selective Bioactive Compounds as an Angiotensin Converting Enzyme Inhibitor

**DOI:** 10.1101/2020.08.17.254359

**Authors:** Huma Khan, Tahir Husain, Monika Kataria, Amit Seth, Md. Zubbair Malik, Ashoutosh Dash, Subhash Chand, Mohammad Azhar Khan

## Abstract

Hypertension is one of a major reason of mortality and morbidity and it is associated with heart and renal disease. The aim of this study is to find out the antihypertensive role of bioactive compounds from selected medicinal plants targeting ACE molecule which so far is not known. The plants taken in this study were *Moringa oleifera*, *Azadirachta indica*, and *Hibiscus sabdariffa*. The nitric oxide and superoxide scavenging property vary from 39.50% to 68% and 37.67 % to 75.50 %. respectively. The inhibition of ACE activity was found maximally in methanolic extract of *A. indica* (74 %), followed by H. *sabdariffa* (73.4%), and least in *M. oleifera* (71.8 %). The bioactive chloroform fraction was characterized for the presence of compound using standard techniques such as LCMS and NMR (^13^C-NMR ^1^H-NMR). The results revealed the presence of beta-sitosterol in *M. oleifera*, azadiradionolide in *A. indica* and hibiscitrin in *H. sabdariffa*. The compounds have shown significant low binding energy for hibiscitrin (−12.3kcal/mol), beta-sitosterol (−11.2kcal/mol) and azadiradionolide (−11.3kcal/mol) indicating the high efficacy of binding on the enzyme. While, binding energy of drug captopril was −5.6kcal/mol & enalpril - 8.1kcal/mol in the same pocket of the ACE molecule. Upon subjecting molecular dynamic simulation results indicated that beta sitosterol complex provided more compactness than the hibiscitrin and azadiradionolide compounds. The current study delivers a new perspective for the drug development against systolic blood pressure regulation and also opens new horizons for considering alternate highly potent drug target for hypertension.

## Introduction

Hypertension or high blood pressure is acknowledged as one of the leading factors for human illness and impermanence. This is a persistent medical condition and is the utmost common life threatening non-communicable disease (Gavras, 2009). With the rise of urbanization, there is a change in lifestyles such as smoking, lack of physical activity, unhealthy diet, alcohol, which are major risk factors of hypertension (Alwan, 2010). Many antihypertensive agents, which include potassium-sparing diuretics, central α2-adrenergic agonists, peripheral α1-adrenergic antagonists, central/peripheral adrenergic neuronal-blocking agents, diuretics, β-blockers, calcium-channel blockers, and blockers of the rennin-angiotensin system, such as angiotensin-converting enzyme inhibitors and angiotensin II type 1 receptor blockers are used alone or in combination to treat hypertension. But antihypertensive drugs have many side-effects such as dry cough, muscle cramps, dizziness, extreme tiredness, dehydration, blurred vision, abnormal heart rate, skin rash, hence, the management of hypertension by herbal medicine is an alternative approach (Vardanyan and Hruby, 2017; McLaughlin *et al.*, 1998).

The renin angiotensin system (RAS) is an important regulator of blood pressure and plays a critical role in the pathogenesis of hypertension, renal and cardiovascular disease. The RAS consist of many components including renin, angiotensinogen (AGT), angiotensin converting enzyme (ACE), ACE2, angiotensin subtype2 receptor (AT2R) and angiotensin subtype 1 receptor (AT1R) (Barnas*et al.*, 1997). The ACE (EC 3.4.15.1) converts Ang I to Ang II, and bradykinin to inactive components. The ACE generated potent Ang II peptide excert effect via predominantly activation of AT1R a vasoconstrictor, plays an important role in the pathogenesis of hypertension, suggesting ACE inhibition as a critical target for controlling systolic blood pressure. Consequently, various compounds from medicinal plants had been explored as ACE inhibitors. *Eleusine indica* leaves showed a role in inhibiting ACE activity (Tutor *et al.*, 2018). The *Chrysophyllum cainito* controls blood pressure due to the presence of phenolic compounds (Mao *et al.*, 2015). Quercetin glycoside a predominant bioactive compound present in citrus fruit, buckwheat and onion was reported as a potential ligand of ACE (Muhammad and Fatima, 2015). The occurrence of secondary metabolites i.e. flavonoids, alkaloids, glycosides, lipids and tannins in plant extract reported to play an important role in ACE inhibition (Reddy *et al.*, 2018). Several, *in silico* studies have also suggested that bioactive compounds from medicinal plants and herbs are potential drugs to target the RAS components including ACE (Khan *et al.*, 2019). Therefore, there is a significant scope to explore the natural compounds as a mediator in controlling blood pressure. The selected plants used in the study were based on literature survey on ethnopharmacology of *Moringa oleifera, Hibiscus sabdariffa,* and *Azadirachta indica*. The compounds from *Moringa oleifera* (Niaziminin-A) (Anwar*et al.*, 2007), *H. sabdariffa* (Delphinidin-3-sambubioside) (Ojeda *et al.*, 2010), and *A. indica* (Nimbidin) (Shori and Baba, 2012) were reported to have antihypertensive activity but their mechanism for targeting ACE molecule is unclear. Hence, the aim of this study is to evaluate ACE inhibitory activity of the compounds from these plant extracts.

## Material and Methods

### Sample Collection and preparation

Leaves of *Moringa oleifera, Hibiscus sabdariffa,* and *Azadirachta indica* were collected from the campus of Shoolini University, Solan, Himachal Pradesh (India) and verified with the Botanical Survey of India, Dehradun, Uttrakhand (India) for herbarium No: (271, 272 and 276). The plant leaves were washed and sterilized by mercuric chloride (HgCl_2_) and air dried for 4-5 days. Subsequently, leaves were pulverized by pestle motor and used for extract preparation in methanolic solvent by Soxhlet method. The extract was weighed and stored in a tube,

### Fractionation of the extract

Ten grams of leaves extracts of *Moringa oleifera, Hibiscus sabdariffa,* and *Azadirachta indica* were re-suspended in 300 ml lukewarm distilled water and then transferred into a separating funnel. Chloroform (50 ml x 3 times) was taken in the separating funnel, observed monitored by mild shaking (1-2 min) and the chloroform fraction was collected. Similar process of fractionation was repetited for n-butanol and ethyl acetate solvents to collect fractions of different polarity by different solvents (Matkowski et al., 2008).

### Characterization by LCMS and NMR

The LC-MS analysis was carried out on X-Bridge C18 (2.1 x 50 mm, 3.5 μm) column fitted to an LC-MS 6320 Ion Trap instrument (Agilent Technologies). A gradient of 50 to 100% acetonitrile in 25 mM ammonium acetate buffer was used as a mobile phase and the flow rate was set at 0.4 ml/min. Injection volume was 5μl. ^1^H & ^13^C NMR spectra were recorded on 400 & 100 MHz cryo-magnet spectrometer (Bruker), respectively (Wolfender et al., 2000).

### Nitric oxide (NO) free radical scavenging activity

In nitric oxide assay, 2 ml of 10 mM sodium nitroprusside in 0.5 ml of phosphate buffer saline was mixed with 0.5 ml of plant extract at different concentrations (100-500μg/ml) and the mixture was incubated for 150 min at 23°C. Of this incubating solution, 0.5 ml was added into 1 ml of sulfanilic acid reagent (20% glacial acetic acid) and incubated for 5 min at 23°C. Thereafter, 1 ml of Naphthylethylene diamine dihydrochloride was added and incubated for 30 min at room temperature. Absorbance was read at 540 nm with spectrophotometer (Rao et al., 2016).

The NO radical scavenging activity was calculated according to the following equation: -

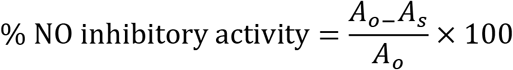

Where Ao = absorbance of the control (blank, without extract); As = absorbance in the presence of the extract.

### Superoxide (O_2_^−^) radical scavenging activity

The superoxide radical scavenging assay was performed as described by Khatua and Acharya, 2016. The 2nM methionine was prepared by dissolving 0.076 gm in 5 ml of distilled water and 0.1 nM EDTA was prepared by 0.004gm in 10 ml of distilled water. Riboflavin was prepared by 0.004 gm in 5 ml of water and 50Mm sodium phosphate buffer of pH 7.8. The reaction was containing 50 μl of sodium phosphate buffer (pH-7.8), 83 μl plant extract in various concentrations (100-500μg/ml), methionine 26μl, EDTA 20μl, Nitrobluetetrazolium 17 μl and riboflavin 4μl. The 96 plate was shaken for 15 seconds and initial absorbance was recorded at 595 nm. The plate was kept under in 15 W light and kept at room temperature for 10 minutes to start reaction. After the incubation the reading was taken at the 595 nm. Ascorbic acid was used as Positive control in different concentration.

The superoxide radical (O_2_^−^) scavenging activity

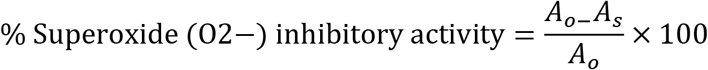

Ao = absorbance of the blank without extract (control); As = absorbance in presence of extract.

### Angiotensin converting enzyme inhibition assay

The inhibition of angiotensin converting enzyme by substrate Hippuryl-histidyl-leucine (HHL) (Sigma-H1635) with benzene sulfonyl chloride and pyridine to convert hippuric acid and histidyl-leucine (Tutor and Chichioco-Hernandez, 2018). The positive control used in this method was captopril.

Forty μl of (100-500μg/ml) plant extract, 20μl of 5mM Hippuryl-histidyl-leucine (Sigma-H1635) and 20μl of angiotensin converting enzyme (Sigma A6678) from rabbit lungs were mixed with 40 μl of sodium borate buffer (pH-8.2) having 0.3 M NaCl. The reaction was start with 15 min pre-incubation of angiotensin converting enzyme with inhibitor at 37 °C for establish contact. After incubation, substrate 20μl of 5mM HHL was further to start Angiotensin converting enzyme reaction. Then incubated mixture aimed at 30 min at 37 °C and then experiment was finished by addition 50 μl of 1m HCl. After this instantaneously by the adding of 100 μl pyridine and 50μl benzenesulfonyl chloride to grow the yellow color. Absorbance was taken at 410 nm using microplate reader

The following formula was used to determine percent inhibition:

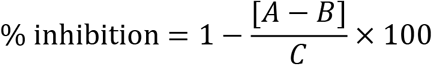

A: Absorbance of Inhibited sample

B: absorbance of Initial sample

C: Absorbance of Uninhibited sample

### *In silico* molecular docking analysis

Crystal structure of C-domain (1O86) with resolution 2 Å of Angiotensin converting enzyme was downloaded from RCSB protein data bank (Corradi HR et al., 2016; Natesh R et al., 2003). The format of the crystal structure of enzyme was changed from .pdb to .pdbqt via AutodockTools-1.5.6. The receptors were further modified by adding polar hydrogens and converting to. pdbqt files used for molecular docking analysis (Muhammad and Fatima, 2015). Chemical structure of isolated phyto-constituents found in methanolic leaf extract of *Moringa oleifera, Hibiscus sabdariffa,* and *Azadirachta indica* were drawn in chemdraw software where 2 D structures were converted into 3 D structure which were further saved as .pdb files (Kashyap et al., 2017). The files were further optimized and converted to. pdbqt files for molecular docking by Auto Dock tools 1.5.6 (Ghersi D and Singh M, 2014). Molecular docking was carried out to evaluate the potential of three compounds against ACE which play an important role in Hypertension. AutoDock Vina achieves a nearly two orders of extent speed-up compared to the other molecular docking software like AutoDock 4 (Hisle et al., 2018; Trott and Olson, 2010). The results of docking are analysed by the association of binding energy with the binding or inhibiting capacity of the ligand for any particular receptor. After docking, the resultant receptor-ligand complexes were generated by UCSF chimera 1.9 which is a highly extensible program for interactive visualization and analysis of molecular structures and docking results to know whether the ligands were bound to active sites of the drug target (Pettersen *et al.*, 2004). The minimum binding energy of bioactive compounds specified potent drug targets for ACE. The best docking compound was selected on the basis of hydrogen bonding with the target. The toxicity of the compounds was predicted by ADME - toxicity (http://lmmd.ecust.edu.cn:8000/predict/). The all isolated compounds i.e. beta-sitosterol, hibiscitrin and azardinolide from *M. oleifera*, *H. sabdariffa*, and *A. indica* respectively were used ADME-toxicity server (Khan *et al.*, 2019).

### Molecular dynamic simulation analysis

All the MD simulations was performed using GROMACS (Wang and Chou, 2009; Sheng-Xiang, 2013) Bio-Simulation package installed in Multi-core enabled Linux Ubuntu system. The force field CHARMM27 (Berendsen, 2005) was used to determine the dynamic behavior and different calculations of proteins (de Souza *et al.*, 1999). The water model SPC216 was used to solvate the system (Angiotensin converting enzyme) and prepare the different using different natural compounds to see the inhibitor of Angiotensin converting enzyme. As a control, MD simulation of 30 ns was carried in water at 300 K. All atoms of Angiotensin converting enzyme and include compounds Beta-sitosterol, Hibiscitrin and Azadiradionolide were equilibrated in cubic box, having size of approximately 73.3 × 73.3 × 73.3 Å. Protein was immersed into water and well equilibrated and overlapping molecules were deleted. Each system was energy minimized with steepest descent, up to a tolerance of 100 kJ mol^−1^Each system was energy minimized with steepest descent, up to a tolerance of 100 kJ mol^−1^nm^−1^ to remove all bad contacts. The overall charge on the system was neutralized with addition of cationic and anionic concentration of Na^+^Cl^−^. To run the simulation, based on the size and 3D spatial orientation of protein, the dimension (x, y and z) of simulation box was defined. All the systems wer prepared in define box, and protein was placed in centre of box, padded around with water and co-solvents. The steepest-descent algorithm along with conjugate gradient was used for the energy minimization process. The system was equilibrated with two ensemble processes, NVT and NPT. Before we start production run, we have to prepared or established the pre-define condition like pH, Temperature. All these information is encapsulated in NVT, NPT and MD parameter files. These physiochemical conditions play a vital role for protein structural and conformational changes. Once, the system is prepared the system is now ready to run.

### Cell toxicity MTT assay

Baby hamster Kidney (BHK-21) cells were cultured at 37°C with 5% CO_2_ in DMEM medium, supplemented with 10% heat inactivated fetal bovine serum. Remove and discard culture medium and then rinse with PBS for removal of extra medium in flask. Briefly rinse the cell layer with trypsin for 2-3 min and incubate in CO_2_ incubator for 2-3 min. Trypsin poured is around 2-3ml and take cells out containing trypsin to observe cells whether they are dispersed (de-adhered) or not. Pour FBS to neutralize the effect of trypsin. Add cell to falcon and centrifuge at 1000 rpm for 5 min to remove debris and get pellet containing cells. Remove supernatant and resuspended pellet in new media. Along the experiments, the cells were monitored by microscopic observation in order to detect any morphological changes. Cells were seeded in triplicate at 10^4^ cells/200 μL per well in a sterile 96-well plate. The cells were grown in an incubator (5 % CO_2_ and 37°C) for 24 hours. After the first incubation, culture medium was replaced by 15 μL of variable concentrations of plant extract (0.12 μg/mL – 0.97 μg/mL). The cells were incubated for the second time for 24 hours. After 24 hours 150ul of cell culture media is removed and replaced by 100ul of DMSO and further incubated for 15 minutes. After 15 minutes reading was taken at 490nm (Srivastava et al., 2009).

## Results and Discussion

### Nitric oxide (NO) free radical scavenging activity and Superoxide (O_2_^−^) radical scavenging activity

The nitric oxide free radical scavenging assay estimation was done by method described by Parul *et al.*, 2013. Among the plant extracts undertaken the *Hibiscus sabdariffa* showed percentage inhibition of 68% (IC_50_ value 317.74 μg/mL) followed by the plant extract of *Azadirachta indica* inhibition of 62% and IC_50_ value (320.7473 μg/mL), and *Moringa oleifera* inhibition of 48.50% (IC_50_ value 350.618 μg/mL), (Table-1). While the ascorbic acid (positive control) showed inhibition of 73.50% (IC_50_ value 302.222 μg/mL) (Figure-1 and Table-1). Based on the free radical scavenging activity the IC_50_ value of *Hibiscus sabdariffa* (317.74 μg/mL) were observed closed to IC_50_ of positive control (302.222 μg/mL). The need of the hour is to harness and explore novel plant sources having appreciable bioactivity against NO radical accumulation and the accompanying inflammatory reaction. In the present work, *Azadirachta indica* and *Moringa oleifera* display higher NO free radical scavenging activity as compared to *Hibiscus sabdariffa.* One of the main enzymes involved in NO production is nitric oxide synthase (NOS) (Ondua et al., 2019). NO scavenging activity of the three plants was evaluated spectrophotometrically at 595 nm. The variations in optical density give a fair assessment of the NO scavenging potency of diverse plant extracts. The phytochemical profile of these plant extracts exhibited the presence of flavanoids and phenolic compounds which are important phytobioactives known for their appreciable radical scavenging potential (Kaurinovic and Vastag, 2019). All the three plants analyzed in the present investigations demonstrate ample NO free radical eradication ability and thus exemplify their intrinsic medicinal potential. Superoxide radicals play a key role in signal transmission at the cellular level (Luis et al., 2018). However, their production beyond a certain threshold in cells has proven to be counterproductive (Basu and Hazra, 2006). This ultimately produces a deleterious effect on biomolecules like proteins, lipids and nucleotides (Engwa, 2018). The most potent superoxide inhibitor proved to be *Moringa oleifera* followed closely by *Azadirachta indica.* These plants were traditionally incorporated in folk medications and hence it becomes imperative to prepare a comprehensive biochemical and antioxidant profile so as to harness their antioxidative stress release potential. These plants also displayed higher radical scavenging activity than the positive control used in these experiments. This further underscores the superoxide radical neutralization effect of these medicinal plants capable enough to prevent cell damage. A wide variety of superoxide scavenging phytoconstituents have been identified and characterized which include mangiferin, naringin, quercitin, myticetin among others (Septiana et al., 2010). The elucidation of optimum concentration of these natural superoxide deradicalizers is a key aspect in their use as efficient therapeutic agents.

**Table 1:** IC_50_ value of Nitric oxide (NO) free radical scavenging activity and Superoxide (O_2_^−^) radical scavenging activity.

| S.NO | Sample | Nitric oxide (NO) free radical scavenging activity (IC<sub>50</sub>) | Superoxide (O<sub>2</sub><sup>-</sup>) radical scavenging activity (IC<sub>50</sub>) |
| --- | --- | --- | --- |
| 1 | *Azadirachta indica* | 352.7473 | 269.7179 |
| 2 | *Moringa oleifera* | 350.618 | 271.5385 |
| 3 | *Hibiscus sabdariffa* | 317.74 | 251.8527 |
| 4 | Ascorbic acid<br>(Positive control) | 302.2222 | 243.3071 |

**Figure 1:**
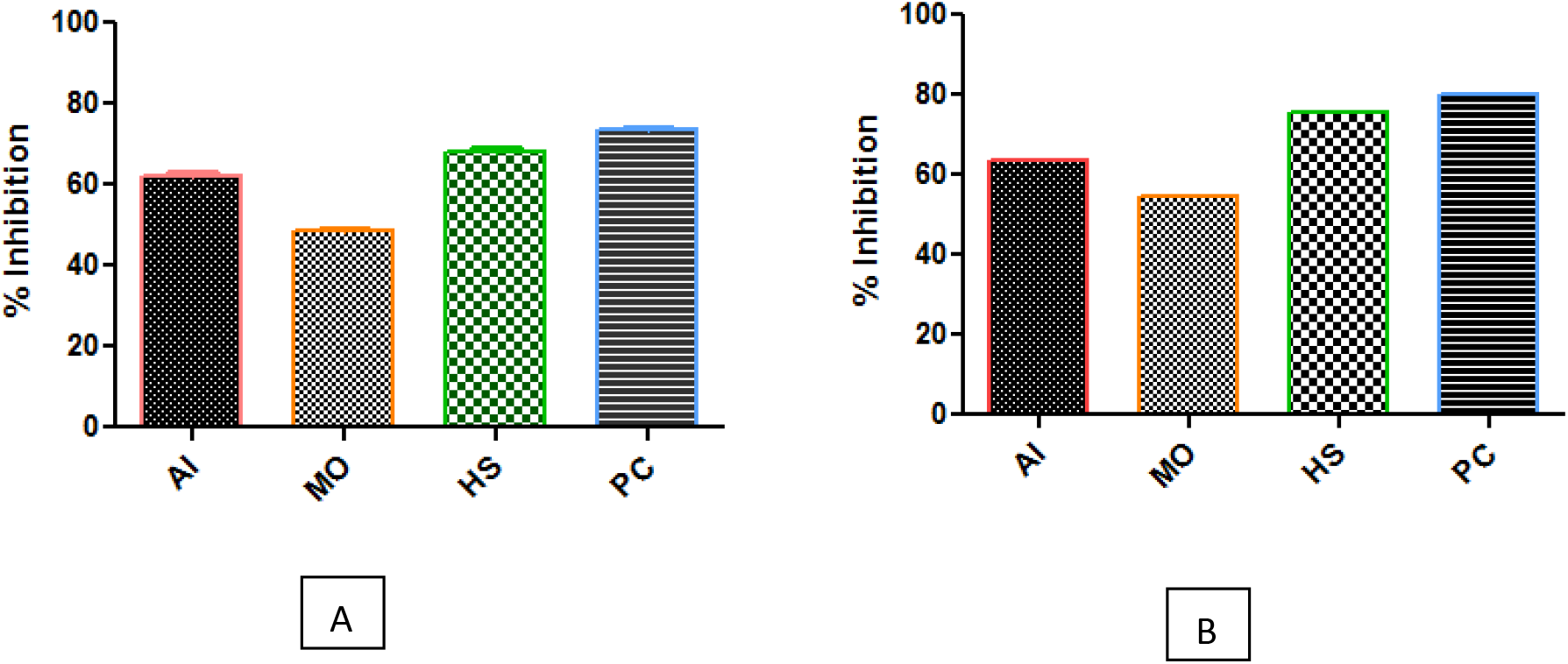
A) percentage inhibition of nitric oxide B) percentage inhibition of SOD.

### LC-MS analysis of active Fractions of *Moringa oleifera, Azadirachta indica,* and *Hibiscus sabdariffa*

While doing the LC-MS analysis of the isolated molecule from chloroform fraction *of Moringa oleifera, we* found a peak of 274 m/z in the mass spectrum (Figure 2). Which was assumed to be a fragmentation of beta-sitosterol 414 m/z. Hence, the primary conformation of the isolated molecule was beta sitosterol (414 m/z). For further conformation we have done ^1^H-NMR and ^13^C-NMR.

**Figure 2:**
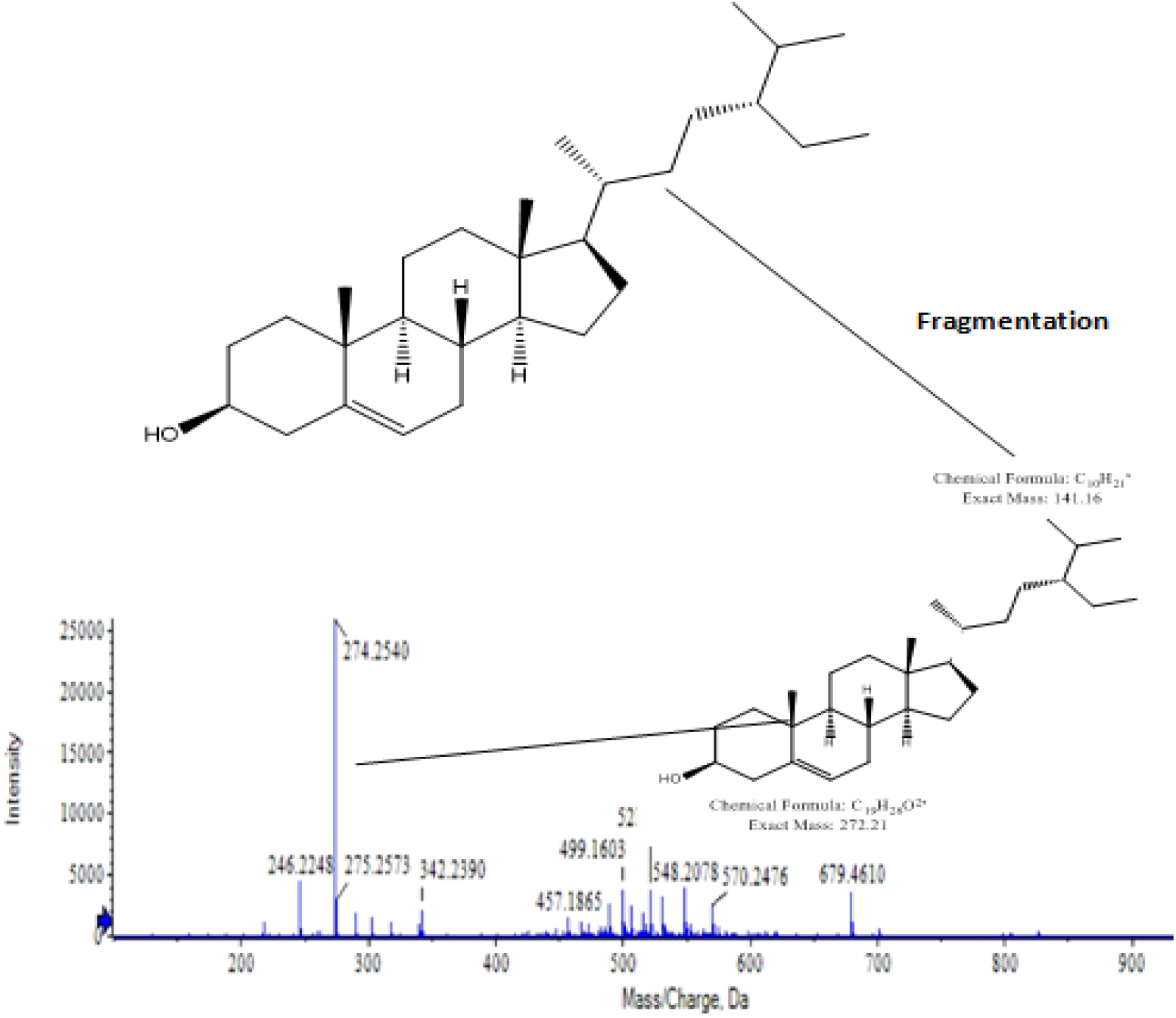
Mass spectrum of different chromatographic peaks of *Moringa oleifera* fraction.

^1^H-NMR and ^13^C-NMR spectra were done using CDCl_3_ as solvent on Bruker Advance II 400 NMR spectrometer at the department of Pharmacy at Guru Jambeshwar University Hisar, Haryana. The ^1^H-NMR spectrum of compound varied between δ 0.5 to 1 ppm. This spectrum shown fifteen protons on carbon C_18_, C_21_, C_26_, C_27_ and C_29_ position in the compound structure (Supplementary Figure -1 and Table-1). proton signal between δ 2.6-4.2 ppm at C_4_, C_7_, C_25_ and 1.2 to 2.5 ppm 47 proton present at different-different carbon position. Chemical shift of δ 5.3 corresponds to oleifinic proton. ^13^CNMR spectrum of Compound (1) shows signal at C_3_ β-hydroxyl group 17.064 and 11.0 for angular methyl carbon atoms for C_19_ and C_18_ respectively. The C_5_, C_6_, C_22_ and C_23_ appeared to be alkene carbons. The value at 17.064 ppm corresponds to angular carbon atom (C_19_). Spectra show twenty-nine carbon signal including six methyls, nine methylenes, eleven methane and three quaternary carbons (Supplementary Figure-2 and Table-2).

While doing the LC-MS analysis of the chloroform fraction of *Azadirachta indica, we* found a peak of 425.19 m/z. Which was assumed to be a fragmentation of Azadiradionolide m/z 480.574g/mol. The *Moringa oleifera* compound i.e. beta sitosterol, similar to cholesterol was also found in fungi, animal and plants. Beta-sitosterol belong to phytosterol which play role in heart disease, antioxidant, diabetes. These are mainly found in the mixture of stigmasterol. The difference between β-sitosterol and stigmasterol is single bond and double bond respectively. Study done by Pierre et al., (2015) found β-sitosterol from *Odontonema strictum.* Hence, the primary conformation of the isolated molecule was Azadiradionolide, which was 425.19 m/z (Figure-3) (274 m/z). The ^1^H NMR spectrum of compound varied between δ 0.5 to 1.75 showed the 15 proton at C_18_ and C_19_ methyl group carbon (supplementary file Figure-3 and Table-3). In δ 2.30 ppm shown 7 protons at C_6_, C_9_, C_11_ and C_12_, δ 3.72 shown 3 protons, δ 3.30-7.55 shown 8 protons at C_17_, C_23_, C_15_, C_2_, C_1_ and C_21_ (supplementary file Figure-4 and Table-4).

**Figure 3:**
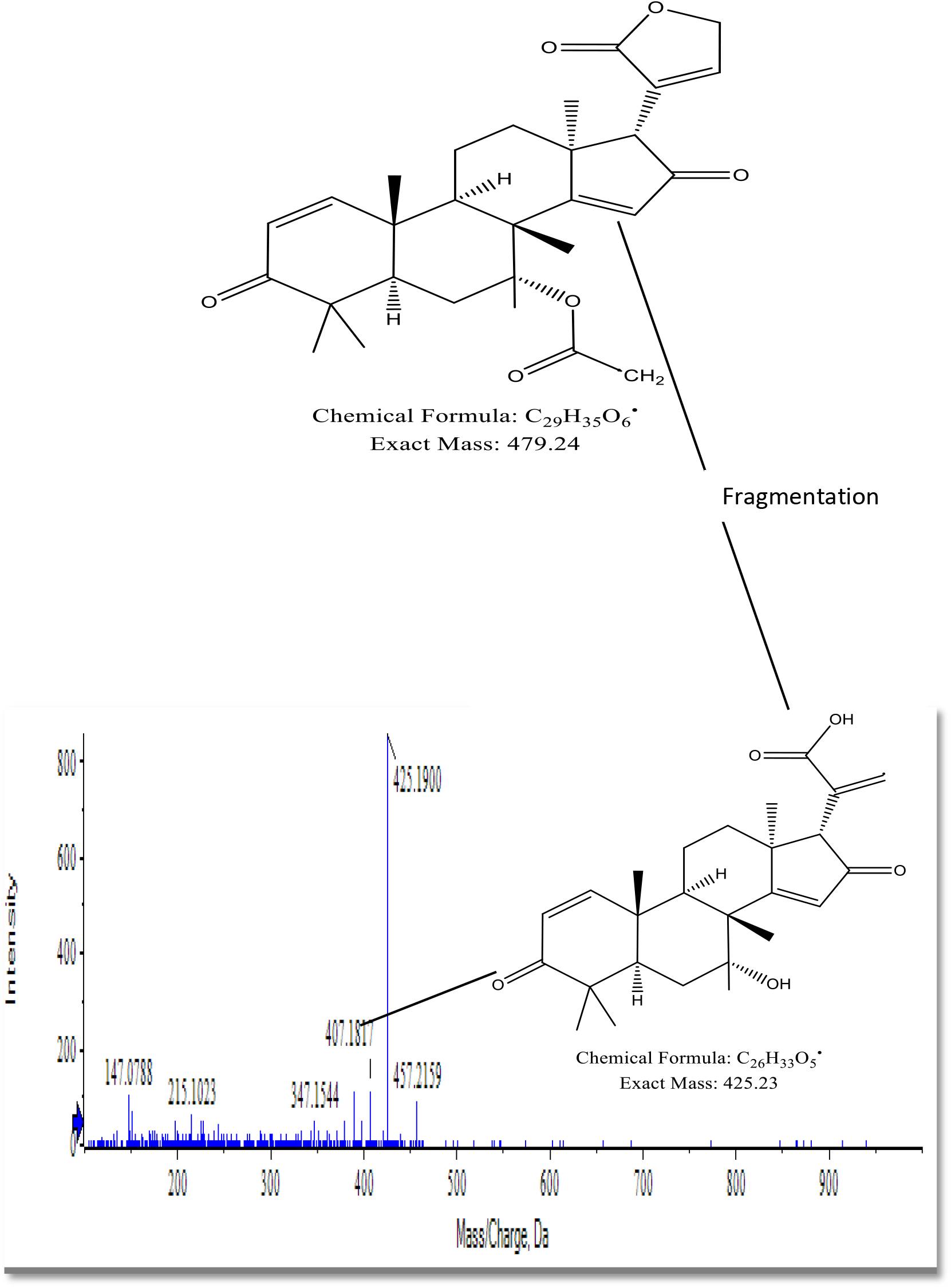
Mass spectrum of different chromatographic peaks of *Azadirachta indica* fraction.

The LC-MS analysis of the chloroform fraction of *Hibiscus sabdariffa, we* found a peak of 274.25 m/z. Which was assumed to be a fragmentation of Hibiscitrin m/z 496.377g/mol (Figure-4). As the molecule was peak of Hibiscitrin after fragmentation is 274.25 m/z (Figure-4). By ^1^HNMR analysis δ 1.01-1.47 ppm shown 12 protons at C_26_, C_27_, C_28_ and C_31_ and δ 2.3-2.8 shown 4 protons at C_23_, C_24_ and C_25_ (supplementary file Figure-5 and Table-5). At δ 3.68-7.28 ppm shown also shown 4 protons at C_13_, C_6_ and C_23_. Methylene group shown at C_28_ (Figure-6 and Table-6).

**Figure 4:**
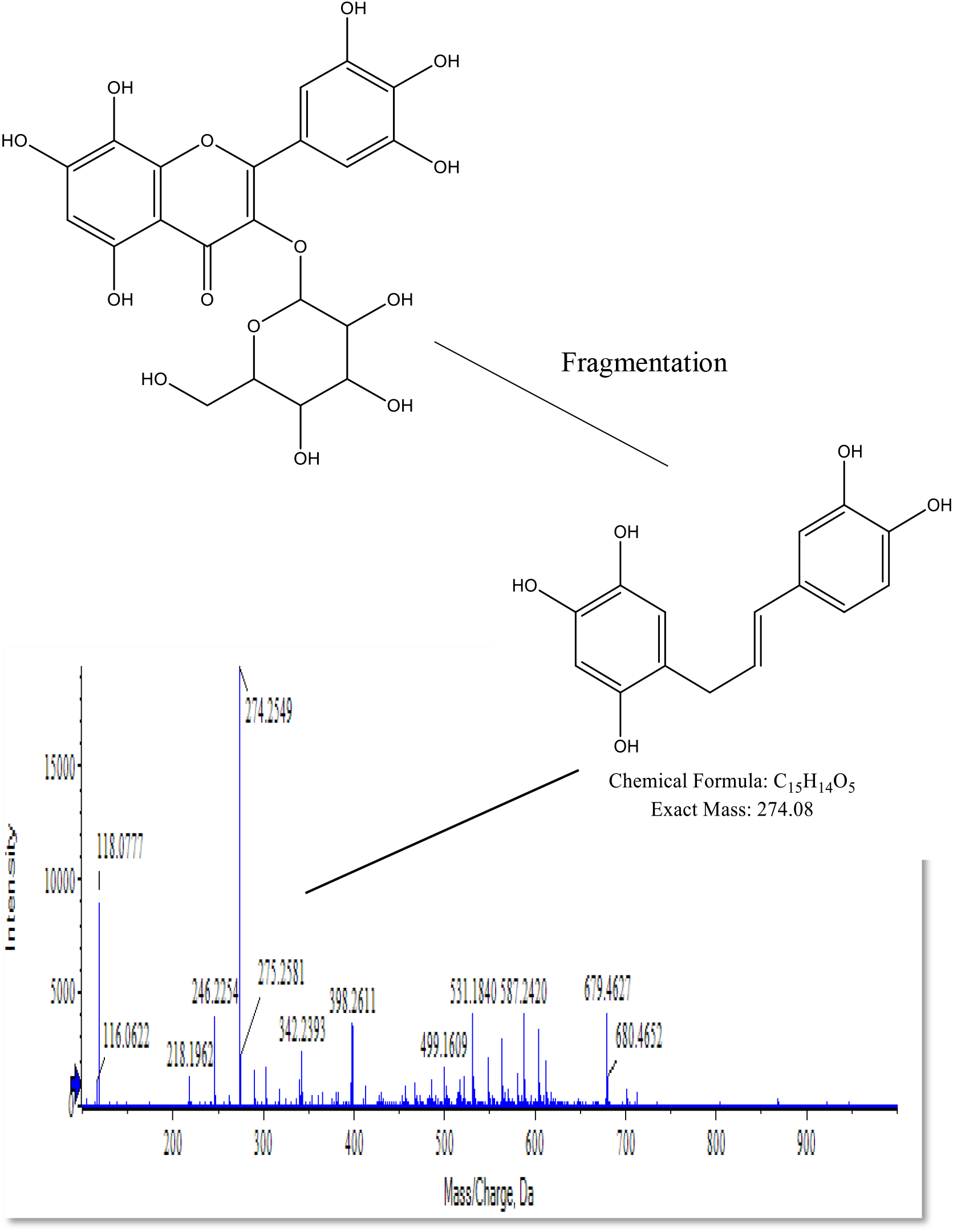
Mass spectrum of different chromatographic peaks of *Hibiscus sabdariffa* fraction.

### Angiotensin converting Enzyme Inhibitory Assay

The six plants extract under this study subjected to ACE inhibitory activity. The percentage inhibition of *Azadirachta indica*, *Hibiscus sabdariffa*, *Moringa oleifera*, *Punica granatum*, and *Allium sativum* were showed 74.00±0.7937 %, 73.47±1.434 %, and 71.80±2.650 %, (Figure-5). The lowest IC_50_ activity were showed by *Azadirachta indica* (255.991 μg/ml) followed by *Hibiscus sabdariffa* (267.722 μg/ml), and *Moringa oleifera* (294.397 μg/ml) (Table-2). Among them based on the IC_50_ the following plants extracts (*Azadirachta indica*, *Hibiscus sabdariffa*, and *Moringa oleifera*) were used for fractionation. The *Azadirachta indica* fraction i.e. n-butanol, ethyl acetate and chloroform showed the percentage inhibition and IC_50_ value were 42.45±1.050% (IC_50_ value 508.5438 μg/mL), 53.25±1.750 % (IC_50_ 373.0502 value μg/mL), and 68.45±0.550 % (IC_50_ value 264.4097 μg/mL) respectively. The *Hibiscus sabdariffa* fraction i.e. n-butanol, ethyl acetate and chloroform showed the percentage inhibition and IC_50_ value were 28.67±1.167 % (IC_50_ value 550.3377 μg/mL), 47.17±1.424 % (IC_50_ value 340.875 μg/mL), and 62.63±1.660 % (IC_50_ value 304.246 μg/mL) respectively. The *Moringa oleifera* fraction i.e. n-butanol, ethyl acetate and chloroform showed the percentage inhibition and IC_50_ value were 32.00±0.981 % (IC_50_ value 461.9108 μg/mL), 43.57±0.263 % (IC_50_ value 338.2692 μg/mL), and 58.50±0.458 % (IC_50_ value 296. 7497 μg/mL) respectively. The percentage inhibition and IC_50_ of positive control (captopril) was 81.13±0.753% (IC_50_ value 203.2158 μg/mL) (Figure-5). Among those fractions the chloroform fractions showed best inhibition activity. Tutor JT and Chichioco-Hernandez CL, 2018 worked on plant extract and ethyl acetate fraction of *Eleusine indica* and calculated percentage inhibition is 68.84% and 51.51% respectively. In our study n-butanol, ethyl acetate, chloroform fraction and extract of *A. indica* shows 30.8%, 45.5%, 63.9% and 67.0% inhibition respectively. The positive control captopril shows 76.0% inhibition for ACE. Deo et al., 2016 and Khan et al., 2019 reported that the 50% inhibition of enzyme by plant extract and fraction consider is active.

**Figure 5:**
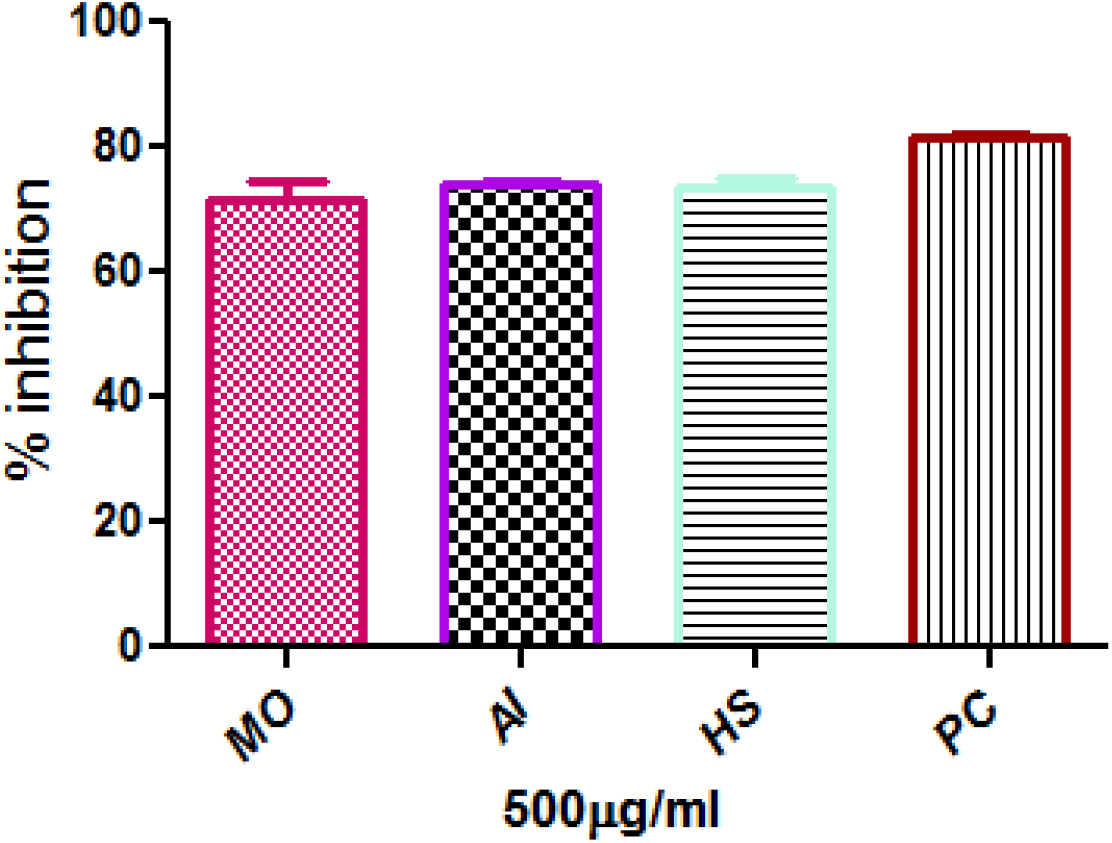
Percentage inhibition of *Moringa oleifera* MO, *Azadirachta indica* (AI), *Hibiscus sabdariffa* (HS), and Positive control (PC)

**Table 2:** IC50 value of plant extract.

| S.No | Plants | IC<sub>50</sub> µg/mL |
| --- | --- | --- |
| 1 | *Moringa oleifera* | 294.397 |
| 2 | *Hibiscus sabdariffa* | 267.722 |
| 3 | *Azadirachta indica* | 255.991 |

ACE is a significant enzyme playing a critical role in regulation of blood pressure. Various plant constituents have the ability to inhibit the action of ACE. The active site of ACE is influenced by the phytochemicals present in neem plant. Flavonoids, terpenoids and phenolic compounds have the ability to forge chemical bonds with the amino acids present in close proximity with the active centre of ACE. These new chemical interactins tend to produce distortions in the catalytic centre of ACE thus causing inhibition. Since ACE is a metalloenzyme having zinc atom at its active site, a majority of hydrogen bonds and chemical bridges involve not just conserved amino acids of the active site but also zinc atom (Yusuf et al., 2000). In this study the ACE inhibiting compound is identified as azadiradionolide. It is a terpenoid comprising multiple rings. The tetracyclic nature of azadiradionolide is responsible for high ACE inhibitory activity of neem extract. The presence of carbonyl, lactones and methyl groups in azadiradionolide is responsible for enhancing the potency of neem plant extract against ACE. These chemical groups may be acting as causative chelating agents for the zinc atom located at the nerve centre of ACE. The in silico studies reveal that azadiradionolide due to its unique structure is able to gain access into the active site of ACE, establish chemical interactions simultaneously with amino acids and zinc atom and causing alterations in the 3-D structure of catalytic protein thus proving to be a successful ACE inhibitor.

### *In silico* study of antihypertensive bioactive compounds against Renin angiotensin components

The *in silico* study is based on the lock and key theory for drug design. This is used for check the biological activity of the natural and synthetic compound and it is time consuming, reasonably and accurate method. The docking of the ligand and enzyme was done by autodock vina software and visualize the hydrogen bonding by chimera and PyMOL. The hydrogen bonding of enzyme and ligand is the important aspect for ligand and enzyme complex formation. In the present study natural isolated compounds targeted the ACE enzyme which has a role in blood pressure regulation. The Angiotensin-converting enzyme converts angiotensin I to angiotensin II a potent peptide hormone exerts its effect predominantly via AT1R (Donoghue et al., 2000). The bioactive molecule reduces the action of angiotensin converting enzyme and control the systolic blood pressure. The fractions of the selected plants were subjected to LCMS and ^1^H-NMR and ^13^C-NMR and following compounds were analyzed. Beta-sitosterol was determined in chloroform fraction of *Moringa oleifera*, Azadiradionolide was determined in chloroform fraction of *Azadirachta indica* and Hibiscitrin found in chloroform fraction of *Hibiscus sabdariffa*. All three compounds were docked with one of the major cascade renin angiotensin component i.e. angiotensin converting enzyme. The compounds i.e Hibiscitrin, beta-sitosterol, and azadiradionolide were showed very low binding energy i.e. −12.3kcal/mol, −11.2kcal/mol and −11.3kcal/mol binding energy with ACE crystal structure respectively. The standard drugs are used in this study is captopril and enalpril which shown −5.6kcal/mol and −8.1kcal/mol binding energy with ACE. The Hibiscitrin compound was interacting with six amino acid residues i.e. Asn-263, Gly-254, Thr-358, His-331, Tyr-498, Lys-489 (Table-3, Table-4). This Hibiscitrin compound shown hydrophobic interaction with ten amino acid residues i.e. His 491, Ala322, Gln 259, Asp255, Thr 144, Glu 262, Glu 431, Ser 357, Phe 435, Phe 505. The beta-sitosterol shown hydrophilic interaction with Asn 145, Gly 254 and Thr 144 and hydrophobic interaction with Thr 352, Asp 255, Asp 340, Leu 139, His 331, Gln 355, and Asp 354 amino acids residues. The compound isolated from neem i.e. azadiradionolide has shown hydrophilic interaction with Asn 145, Asn 263, Thr 144 and hydrophobic interaction Asp 354, Gly 254, Ala 148, Asp 255, Leu 147, Leu 139, Asp 140, His 331, Gln 355, and Thr 352 amino acid residues. It can be observed that compounds Hibiscitrin a greater number of hydrogen bond interactions with various amino acid residues of the Angiotensin converting enzyme residues. We believe that these hydrogen bond interactions play an important part in the inhibition of the Angiotensin converting enzyme residues, and ultimately better anti-inflammatory activity of the compounds Hibiscitrin (Figure-6). An overlay of the docked pose of Azadiradionolide and Beta-sitosterol also exhibited that these compounds are superimposable with each other (Figure-7, Figure-8.). This indicates that the active site of the Angiotensin converting enzyme residues can accommodate these compounds, which synergizes the Angiotensin converting enzyme residues inhibitory activity of the compounds Azadiradionolide and Beta-sitosterol. All these three compounds have shown better binding affinity in comparison to standard drugs (captopril and enalpril). Further these compounds are used for toxicity of the molecule by ADMETSAR software. The toxicity of the compound is the main reason of drug failure in the drug development field (Table-5).

**Table 3:** *In silico* study of isolated compounds with angiotensin converting enzyme.

| S.No | Plant name | Chemical compounds | Type of compound | Molecular formula | Molecular weight | Binding affinity |
| --- | --- | --- | --- | --- | --- | --- |
| 1. | *Hibiscus sabdariffa* | Hibiscitrin | Natural | C<sub>21</sub>H<sub>20</sub>O<sub>14</sub> | 496.4g/mol | -12.3 kcal/mol |
| 2. | *Azadirachta indica* | Azadiradionolide | Natural | C<sub>28</sub>H<sub>34</sub>O<sub>6</sub> | 466.6g/mol | -11.3 kcal/mol |
| 3. | *Moringa oleifera* | Beta sitosterol | Natural | C<sub>29</sub>H<sub>50</sub>O | 414.7g/mol | -11.2 kcal/mol |
| 6. | Positive control | Enalapril (standard drug) | Synthetic | C<sub>20</sub>H<sub>28</sub>N<sub>2</sub>O<sub>4</sub> | 376.453g/mol | -8.1kcal/mol |
|  |  | Captopril (standard drug) | Synthetic | C<sub>9</sub>H<sub>15</sub>NO<sub>3</sub>S | 217.283g/mol | -5.6kcal/mol |

**Table 4:** Hydrogen bonding interaction of the compounds with active amino acid residues of angiotensin converting enzyme.

| S.No | Compound | Active residues | Hydrogen Bonding length | Average bond length |
| --- | --- | --- | --- | --- |
| 1 | **Hibiscitrin** | TYR (498)<br>LYS (489)<br>HIS (263)<br>TYR (358)<br>SER (260)<br>ASN (263)<br>GLY (254) | 3.21<br>3.24<br>3.20<br>3.09<br>2.92<br>2.95<br>2.71 | 2.9 |
| 2 | **Azadiradionolide** | THR (144)<br>ASN (263)<br>ASN (145) | 3.00<br>2.80<br>3.09 | 2.6 |
| 3 | **Beta sitosterol** | TYR (372),<br>ASP (366),<br>HIS (388),<br>GLY (382) | 2.70<br>3.01<br>3.09<br>2.74 | 2.8 |
| 13 | **Captopril** | ARG (522)<br>GLU (411) | 1.75<br>2.85 | 2.1 |
| 14 | **Enalapril** | TYR (523)<br>HIS (353) | 2.55<br>2.87 | 2.7 |

**Figure 6:**
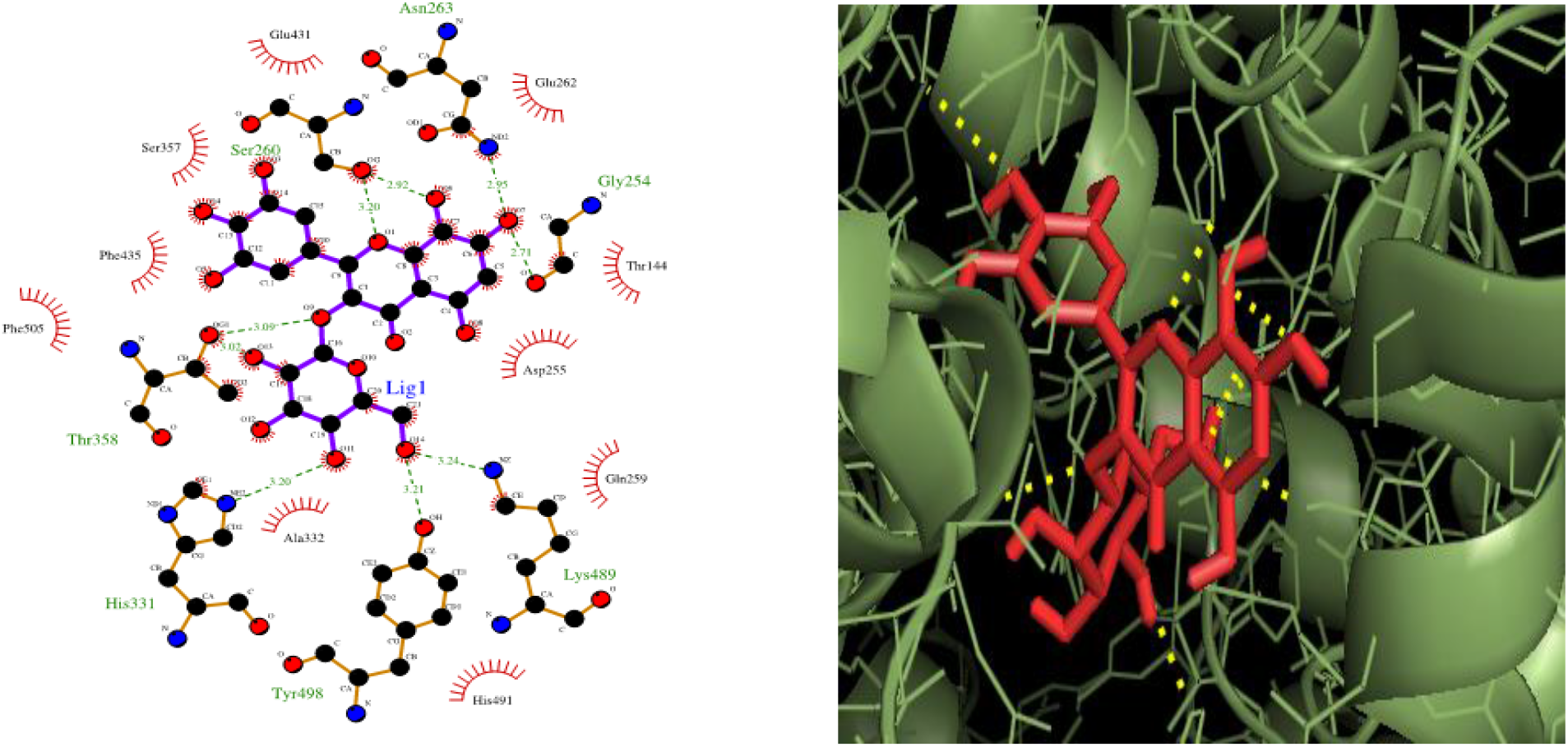
Ligplot of Hibiscitrin ligand from *Hibiscus sabdariffa* fraction with active amino acid residue of ACE.

**Figure 7:**
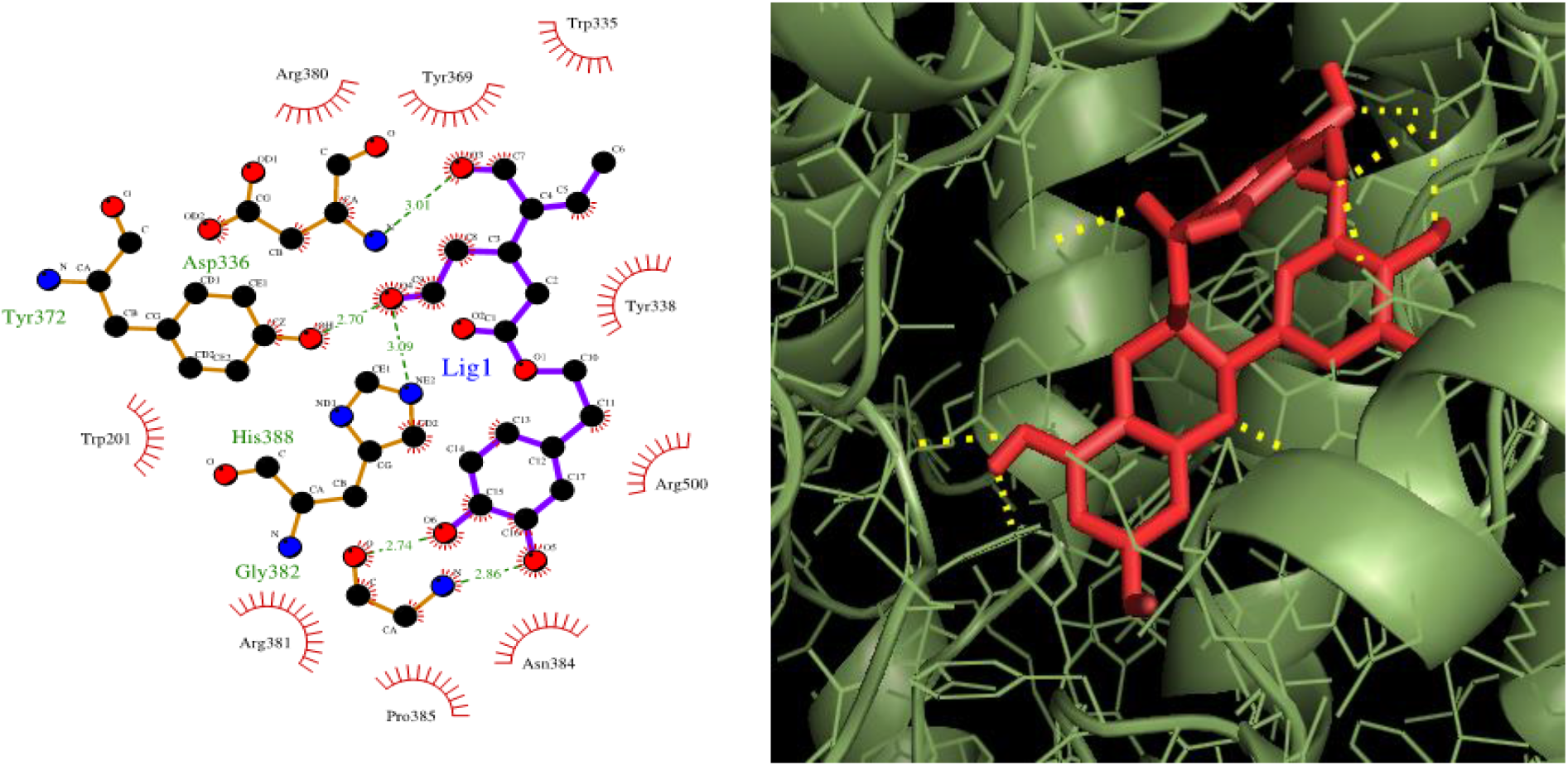
Ligplot of beta sitosterol ligand from *Moringa oleifera* fraction with active amino acid residue of ACE.

**Figure 8:**
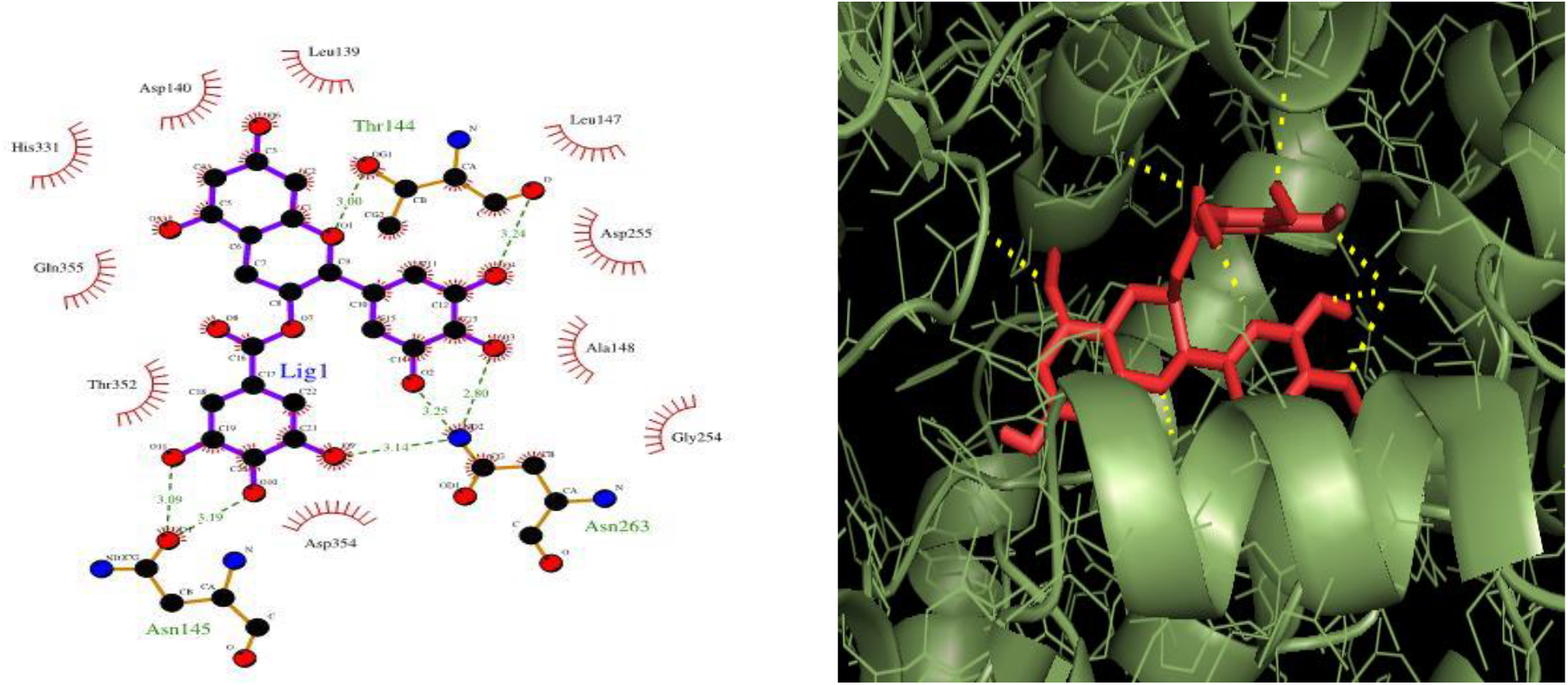
Ligplot of Azadiradionolide ligand from *Azadirachta indica* fraction with active amino acid residue of ACE.

**Table 5:** Toxicity Profile of the Potential Compounds (WI: Weak inhibitor, NI: Non-inhibitor, NT: Non-toxic, T: Toxic NC: Non-carcinogen, C: Carcinogen, HT: High toxic, LT: Low toxic, NRB: Not readily biodegradable)

| Parameters<br>↓<br>Compounds | Human Ether-a-gogo- Related Gene Inhibition | AMES Toxicity | Carcinogens | Fish Toxicity | Tetrahymena Pyriformis Toxicity | Honey Bee Toxicity | Acute Oral Toxicity |
| --- | --- | --- | --- | --- | --- | --- | --- |
| Beta-sitosterol | WI | NT | NC | HT | HT | HT | IV |
| Hibiscitrin | WI | NT | NC | HT | HT | HT | I |
| Azadiradionolide | WI | NT | NC | HT | HT | HT | III |

### Molecular Dynamic Simulation Study

MD simulation study has been also executed to investigate the conformational stability of Beta-sitosterol, Hibiscitrin and Azadiradionolide in active site of Angiotensin converting enzyme in water at 300 K. To examine the structural stability of Angiotensin converting enzyme and complex with beta-sitosterol, hibiscitrin and Azadiradionolide. We monitored the time evolution plot of all *C*_*α*_atom RMSD, Rg, RMSF and SASA. In the fourth parameter the angiotensin converting enzyme with beta sitosterol complex has shown a reduced SASA from 310 nm^2^ to 285 nm^2^ up to 5 ns. The solvent accessibility surface area for the of angiotensin converting with beta sitosterol complex indicated a more compact size at the end of the simulation period. SASA. In this case of hibiscitrin (depicted in green), was reduced from 310 nm^2^ to 290 nm^2^ and in the case of azadiradionolide reduced from 310 nm^2^ to 280 nm^2^. This indicates that beta sitosterol complex will provide more compactness than the hibiscitrin and azadiradionolide, and aromatic ring has less exposed the later on complex. We have detected that complex of protein and compounds shows relatively higher SASA value than wild type protein this could be explain by the presence of relatively larger hydrophobic packed core region in native protein as compare to protein compound complex. In one such similar studies these parameters (RMSD, Rg, SASA and RMSF) were studied by Fang *et al.*, (2019) for ACE that support that support our study.

### Radius of gyration

Another important parameter, radius of gyration (R_g_) is used to determine the dynamic adaptability of A CE in water and three different compounds which namely, beta sitosterol, hibiscitrin and azadiradionolide. A time evolution R_g plot_ of backbone atoms (Figure-9 A) shows that the native conformation of ACE is retained in water R_g_ trajectory in water (Black) quickly achieves the stable equilibrium in few nanoseconds. The average R_g_ values of 2.37±0.01 nm is maintained throughout the simulation period. In the presence of beta sitosterol (depicted in red), 298 K temperature, the system is decreased in the fluctuation of trajectory is suggested that initial compaction in structure with comparison to water (depict in black) an average value is 2.31 ± 0.02. while the previous condition is remaining same and only change the compound instead of Beta sitosterol to used hibiscitrin (depicted in green). Now, the system is gradually increase the fluctuation throughout the simulation relatively less stable than Beta sitosterol, and the average R_g values_ of 2.34 ± 0.04 which indicates a less stable structure and system is expanded. In the presence of Azadiradionolide, 298 K temperature, the Angiotensin converting enzyme is more stabilized than other three compounds. In R_g_ trajectory of Azadiradionolide (Blue line) depicted that quickly achieves stability at 5 ns and system remain stable and the average R_g values_ of 2.28 ± 0.01. nrg for native protein is comparatively more than protein compound complex. The Rg results shows that native protein was more compact after the binding of compound.

**Figure 9:**
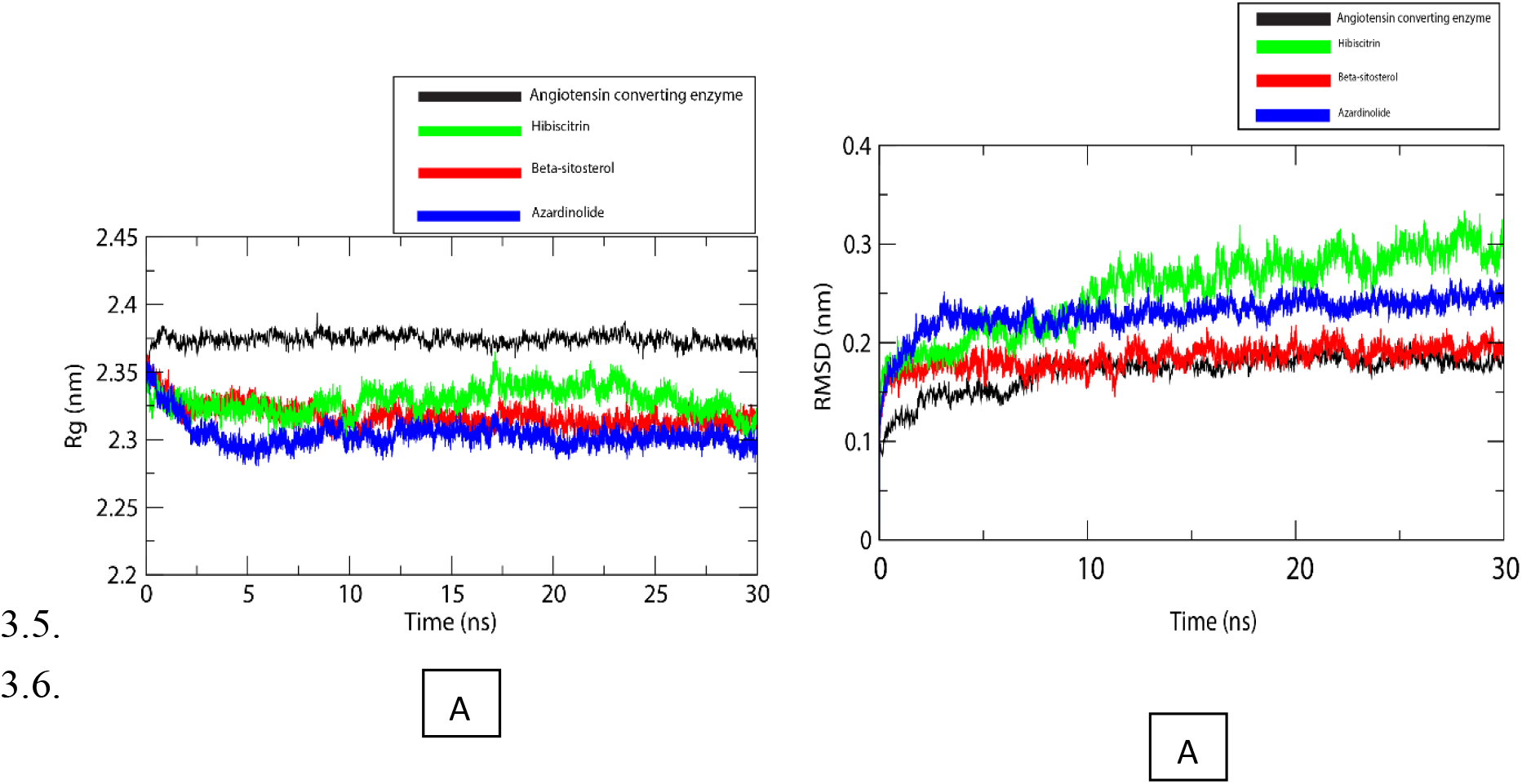
Radius of gyration, RMSD analysis of ACE and bioactive compounds over 30ns simulation.

### Root means square distance (RMSD)

RMSD are calculated as a function of time, with respect to the initial conformation and its shown in Figure-9 B. Visual inspection of this plot show that Angiotensin converting enzyme in water (black) is unstable during the 0-10 ns time interval and after 10-30 ns system have attained the equilibrium is maintained till the end of simulation of 30 ns. Addition of in the presence of Beta sitosterol (depicted in red), 298 K temperature to the system resulted in structural perturbation is slightly come into picture at time 0–10 ns which is relatively more than water and the simulation end up with an increase in RMSD of ~0.012 nm as compare to native condition. The remaining condition is remain intact only compound is change Hibiscitrin (depicted in green) the morphology of structure is change more than Beta sitosterol and the system remain unstable during the entire time evolution of simulation. The presence of azadiradionolide (depicted in blue) at 298 K temperature, the ACE in presence of azadiradionolide the system is unstable during the 0-10 ns time interval and after 10-30 ns system. While the behavior of morphology is in between the Beta sitosterol and Hibiscitrin.

### Root means square fluctuation (RMSF)

To capture the dynamics progression of Angiotensin converting or flexibility of structure in water and three different complex with compounds of the Beta-sitosterol (depicted in red), Hibiscitrin (depicted in green) and are Azadiradionolide (depicted in blue) are depicted in Figure-10 A. RMS fluctuations showed that the most flexible residues 291–301, 338-354 are located in loop regions and 324–347 are located in the loop regions, which connect the Beta-sheets to alpha-helices. RMS fluctuation is more in the of Angiotensin converting +Azadiradionolide (depicted in blue) complex compared to the protein +beta sitosterol (depicted in red) complex. Comparatively, Angiotensin converting +Hibiscitrin (depicted in green) showed high fluctuations at 53–65, 66–144, 145–192 positions, these amino acids mostly constituting the loop region, alpha helices played no role in Azadiradionolide (depicted in blue) binding.

**Figure 10:**
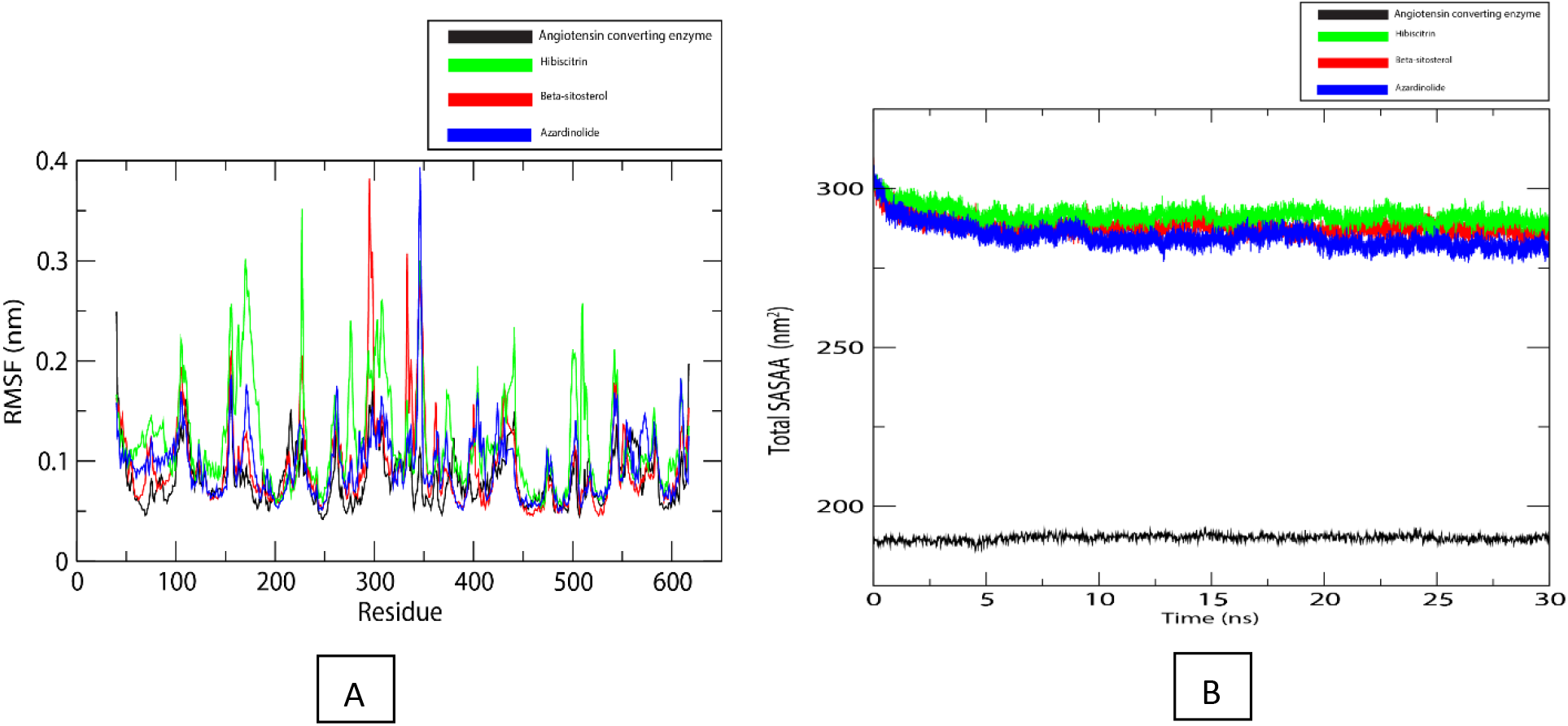
RMSF of ACE with active amino acid residues and SASA of ACE and bioactive compounds over 30ns.

### Solvent accessible surface area (SASA)

The SASA for Angiotensin converting + water (in depicted black colour) was 190nm^2^ at the beginning of the simulation and retained the area throughout the simulation period shown in the Figure 10 B Which is quite stable, apart from this addition of in the presence of Beta sitosterol (depicted in red), 298 K temperature to the system the solvent accessible surface area (SASA) for Angiotensin converting + beta sitosterol was observed to be 310 nm^2^ at the start of simulation; as dynamics proceeded, it continuously decreased to 285 nm^2^. The Angiotensin converting + beta sitosterol complex has shown a reduced SASA from 310 nm^2^ to 285 nm^2^ up to 5 ns.

The solvent accessibility surface area for the of Angiotensin converting + beta sitosterol complex indicated a more compact size at the end of the simulation period. SASA. In this case of Hibiscitrin (depicted in green), was reduced from 310 nm^2^ to 290 nm^2^ and in the case of azadiradionolide reduced from 310 nm^2^ to 280 nm^2^. This indicate that beta sitosterol complex is provide more compactness than the Hibiscitrin and azadiradionolide, and aromatic ring has less exposed the later on complex. We have detected that complex of protein and or compounds shows relatively higher SASA value than wild type protein this could be explain by the presence of relatively larger hydrophobic packed core region in native protein as compare to protein compound complex.

### Cell toxicity assay

To determine the optimum concentration of the fractionated compounds for biological assays the cell toxicity was determined by MTT assay. The chloroform fraction containing predominantly beta-sitosterol, Hibiscitrin and azadiradionolide compounds were subjected to MTT assay on BHK-21 cell lines. The results revealed that with the treatment of bioactive fraction bearing hibiscitrin (HS) compound showed 99.34% viability at 970ug/ml followed by beta sitosterol (BS) with 99.13% viability at 970ug/ml and azadiradionolide (AZ) with 98.67% viability at 970ug/ml. Thus, results indicated that these compounds were least toxic to the cells, further suggesting these compounds are biocompatible and can be utilized as a drug (Figure-11). The Nemudzivhadi and Masoko (2014) showed the 90% cell viability of bioactive compound from *Ricinus communis*. In another study by Raiola *et al.*, (2016) found the 80% cell viability on HEK-293 cell lines by the extract of tomatoes which is lower from the present study. In support to our findings Arevalo *et al.*, (2018) found 50% cell viability by the extract of *M. oleifera* and *A. indica on* MDCK cell lines.

**Figure 11:**
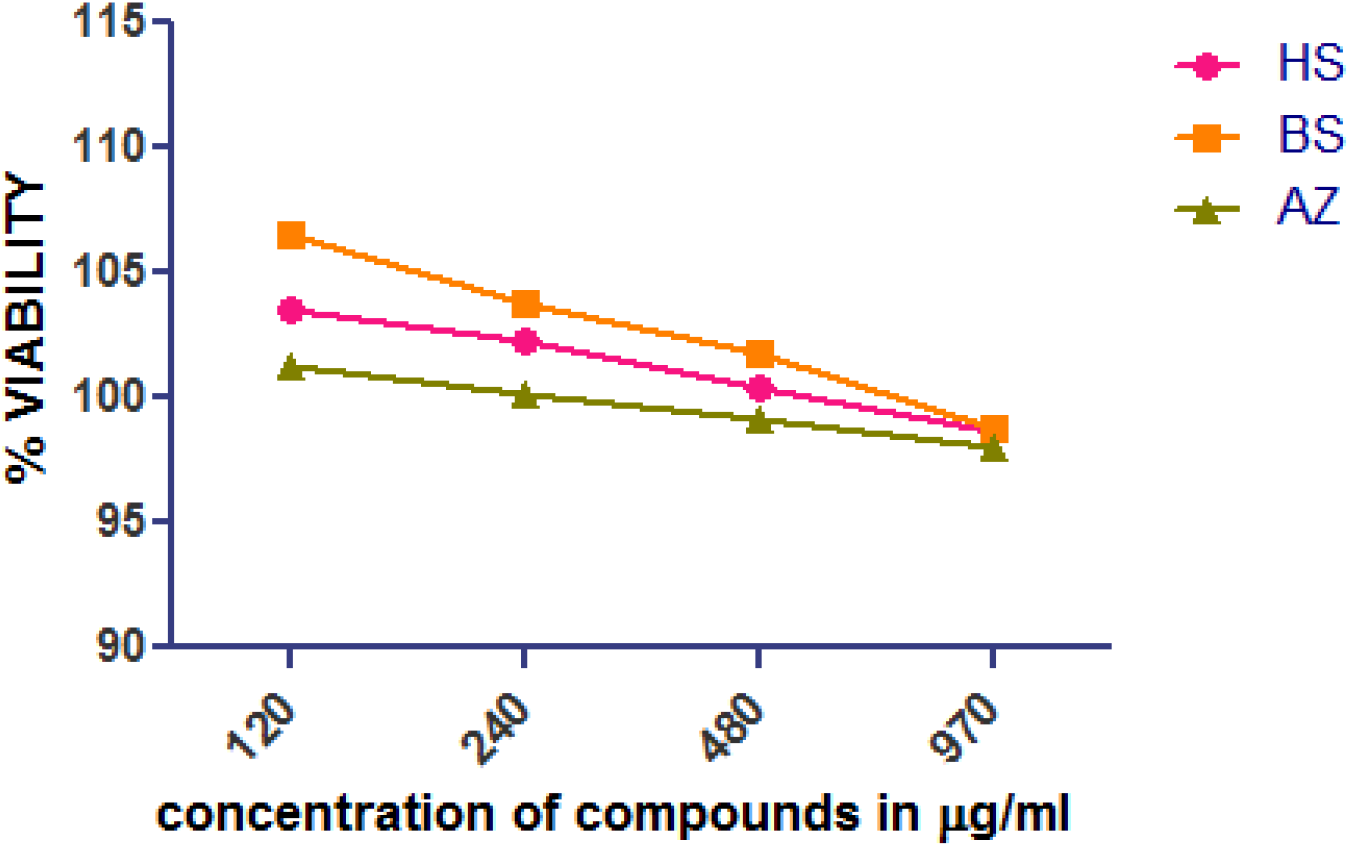
Percentage viability of the compounds Hibiscitrin (HS) beta-sitosterol (BS), and Azadiradionolide (AZ)

### Summary

The present study suggested the consideration of medicinal plants of Himachal Pradesh for targeting the role in hypertension. The isolated and characterized bioactive molecule from *Moringa oleifera, Hibiscus sabdariffa* and *Azadirachta indica* were found to be beta-sitosterol, hibiscitrin and azadiradionolide respectively. The most potent molecule to inhibit the ACE of these was hibiscitrin. So this compound could play a significant role in inhibiting the conversion of angiotensin I to angiotensin II to control the systolic blood pressure. The current study delivers a new perspective for the drug development against systolic blood pressure regulation and also opens new horizons for considering alternate highly potent drug target for hypertension.

## Supplementary file

**Figure 1:**
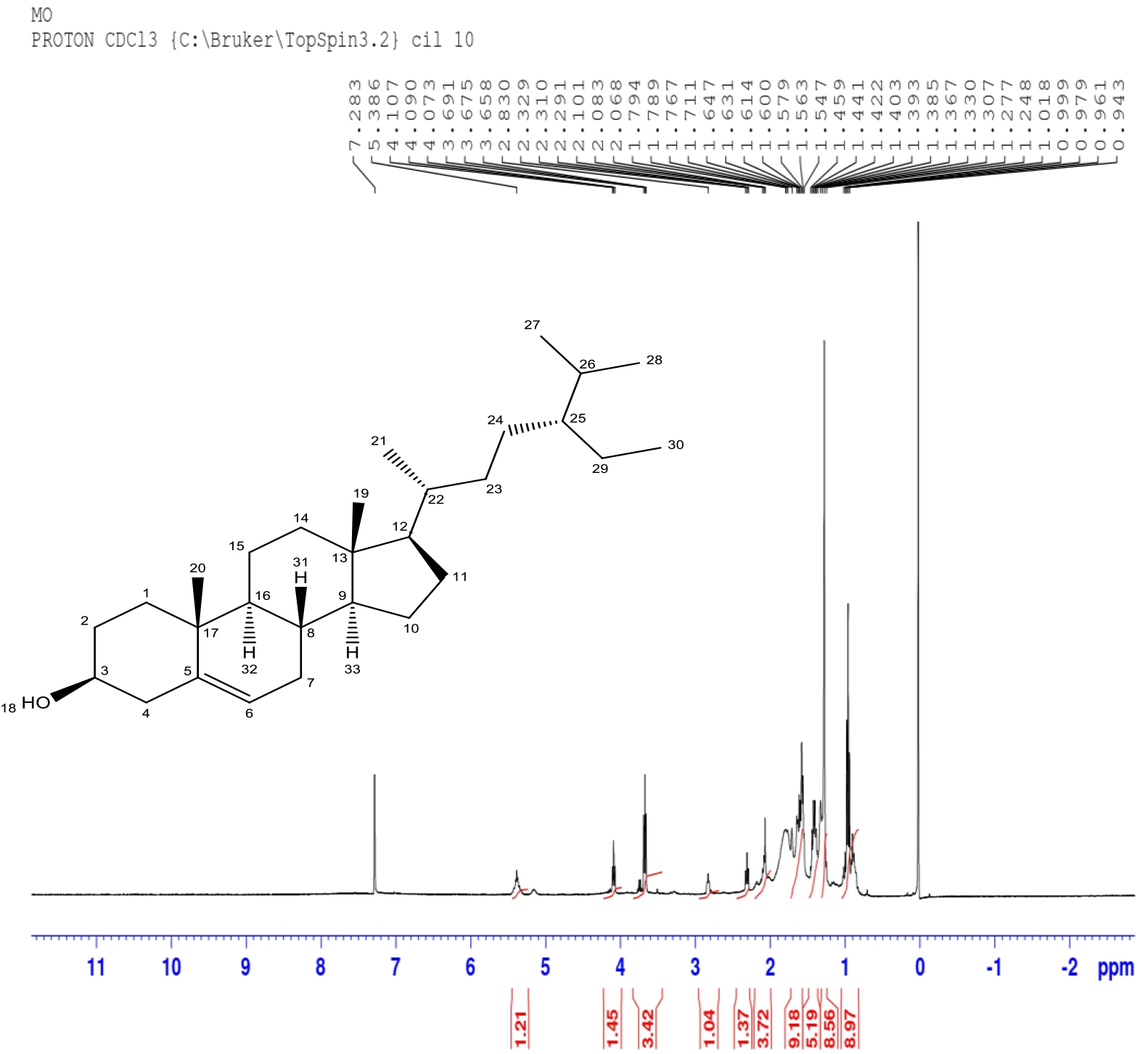
^H^NMR spectrum of different chromatographic peaks of *Moringa oleifera* fraction.

**Table 1:**
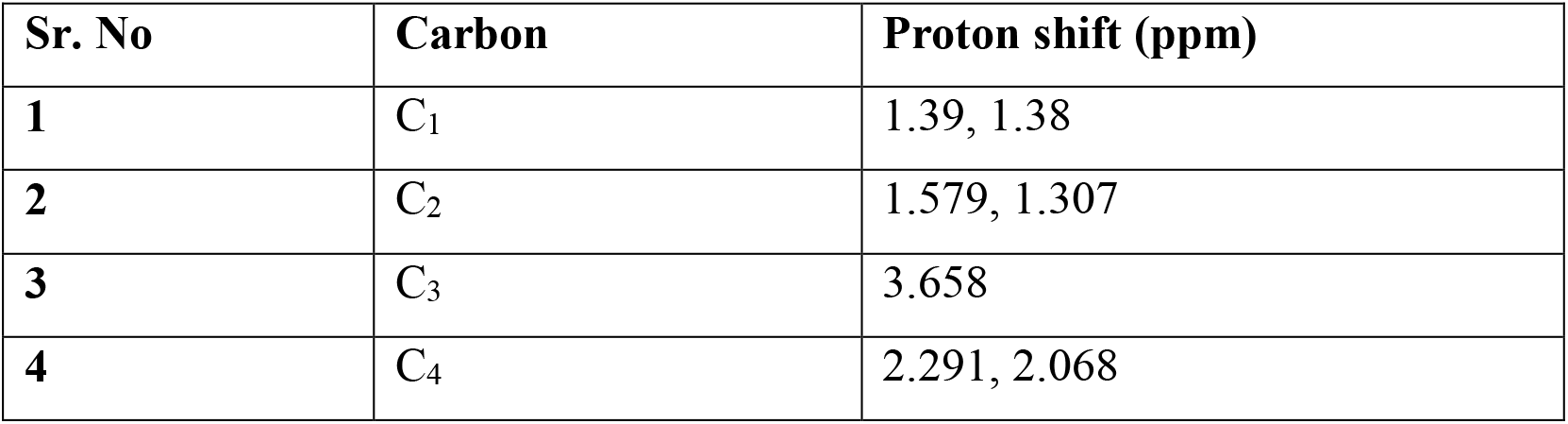
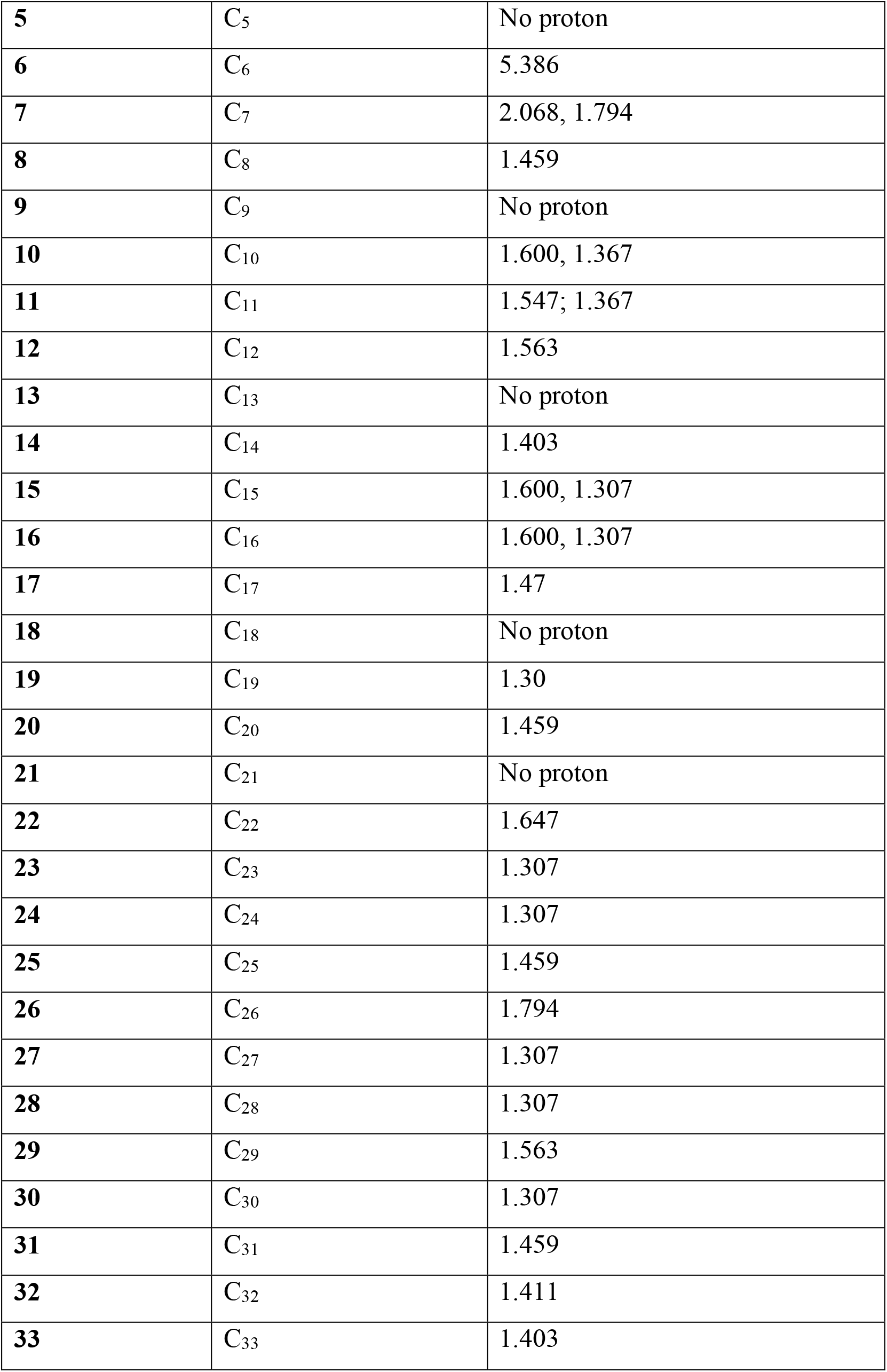
Proton shift position on carbon atom of *Moringa oleifera* fraction.

**Figure 2:**
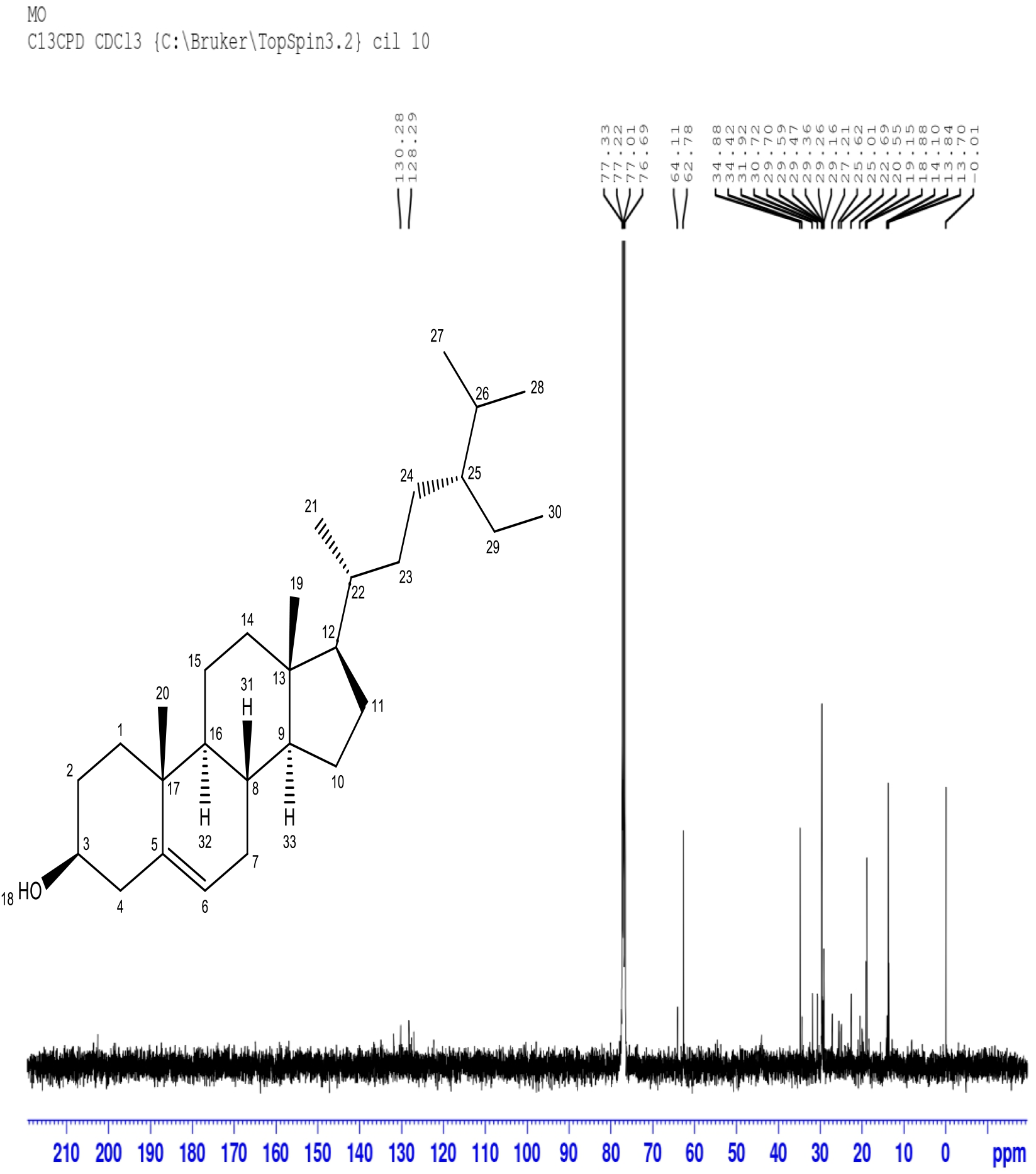
^C^NMR spectrum of different chromatographic peaks of *Moringa oleifera* fraction.

**Table 2:** Carbon shift position on carbon atom of *Moringa oleifera* fraction.

| **Sr. No.** | **Carbon** | **Carbon shift (ppm)** |
| --- | --- | --- |
| **1** | C<sub>1</sub> | 34.88 |
| **2** | C<sub>2</sub> | 31.92 |
| **3** | C<sub>3</sub> | 64.11 |
| **4** | C<sub>4</sub> | 34.88 |
| **5** | C<sub>5</sub> | 34.88 |
| **6** | C<sub>6</sub> | 64.11 |
| **7** | C<sub>7</sub> | 31.92 |
| **8** | C<sub>8</sub> | 31.92 |
| **9** | C<sub>9</sub> | 62.78 |
| **10** | C<sub>10</sub> | 25.62 |
| **11** | C<sub>11</sub> | 25.62 |
| **12** | C<sub>12</sub> | 62.78 |
| **13** | C<sub>13</sub> | 34.88 |
| **14** | C<sub>14</sub> | 34.88 |
| **15** | C<sub>15</sub> | 20.55 |
| **16** | C<sub>16</sub> | 62.78 |
| **17** | C<sub>17</sub> | 34.88 |
| **18** | C<sub>18</sub> | No carbon |
| **19** | C<sub>19</sub> | 13.70 |
| **20** | C<sub>20</sub> | 19.3 |
| **21** | C<sub>21</sub> | 19.3 |
| **22** | C<sub>22</sub> | 34.88 |
| **23** | C<sub>23</sub> | 34.42 |
| **24** | C<sub>24</sub> | 22.69 |
| **25** | C<sub>25</sub> | 34.88 |
| **26** | C<sub>26</sub> | 30.72 |
| **27** | C<sub>27</sub> | 20.55 |
| **28** | C<sub>28</sub> | 22.69 |
| **29** | C<sub>29</sub> | 22.69 |
| **30** | C<sub>30</sub> | 13.70 |
| **31** | C<sub>31</sub> | No carbon |
| **32** | C<sub>32</sub> | No carbon |
| **33** | C<sub>33</sub> | No carbon |

**Figure 3:**
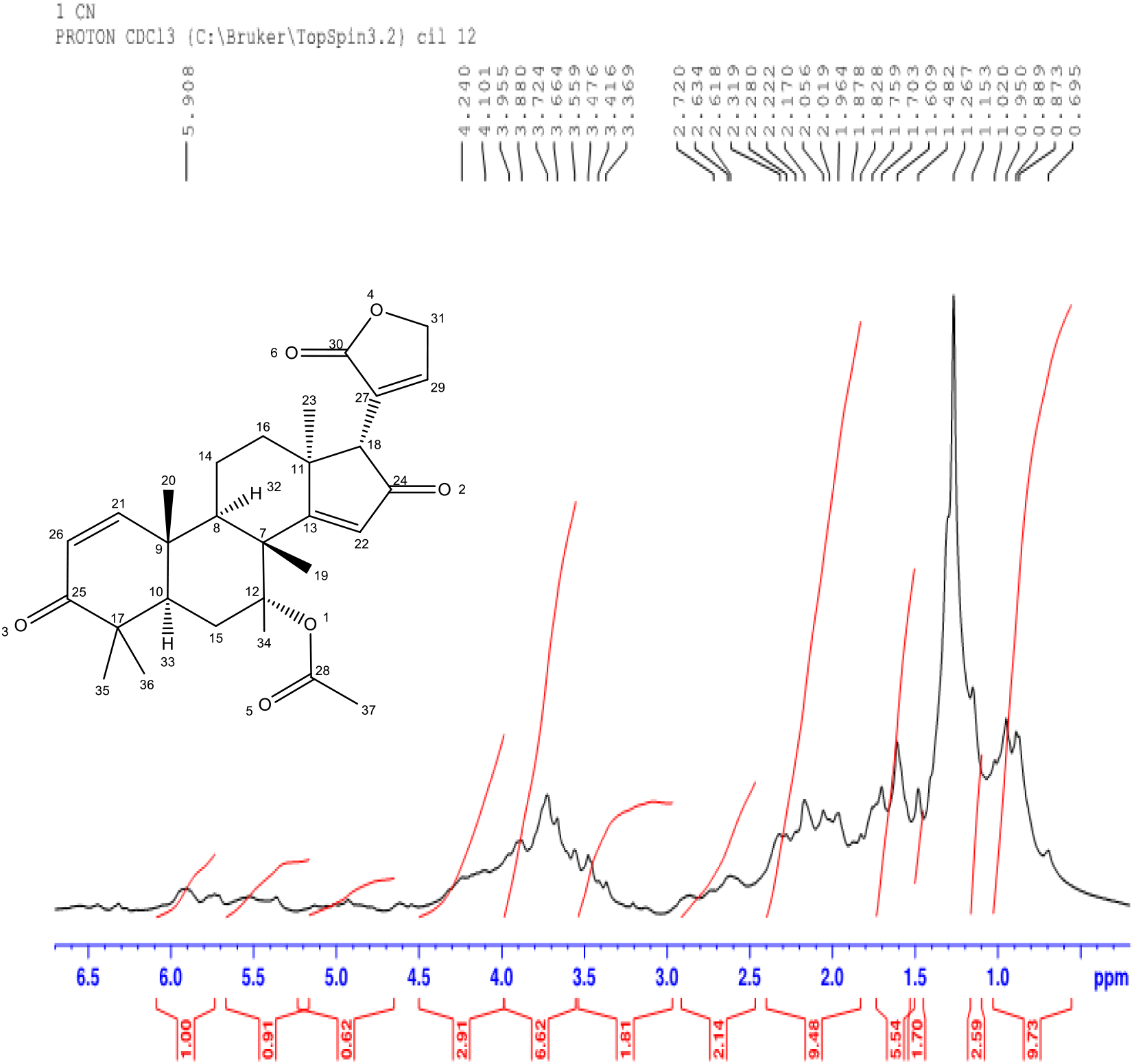
^H^NMR spectrum of different chromatographic peaks of *Azadirachta indica* fraction.

**Table 3:**
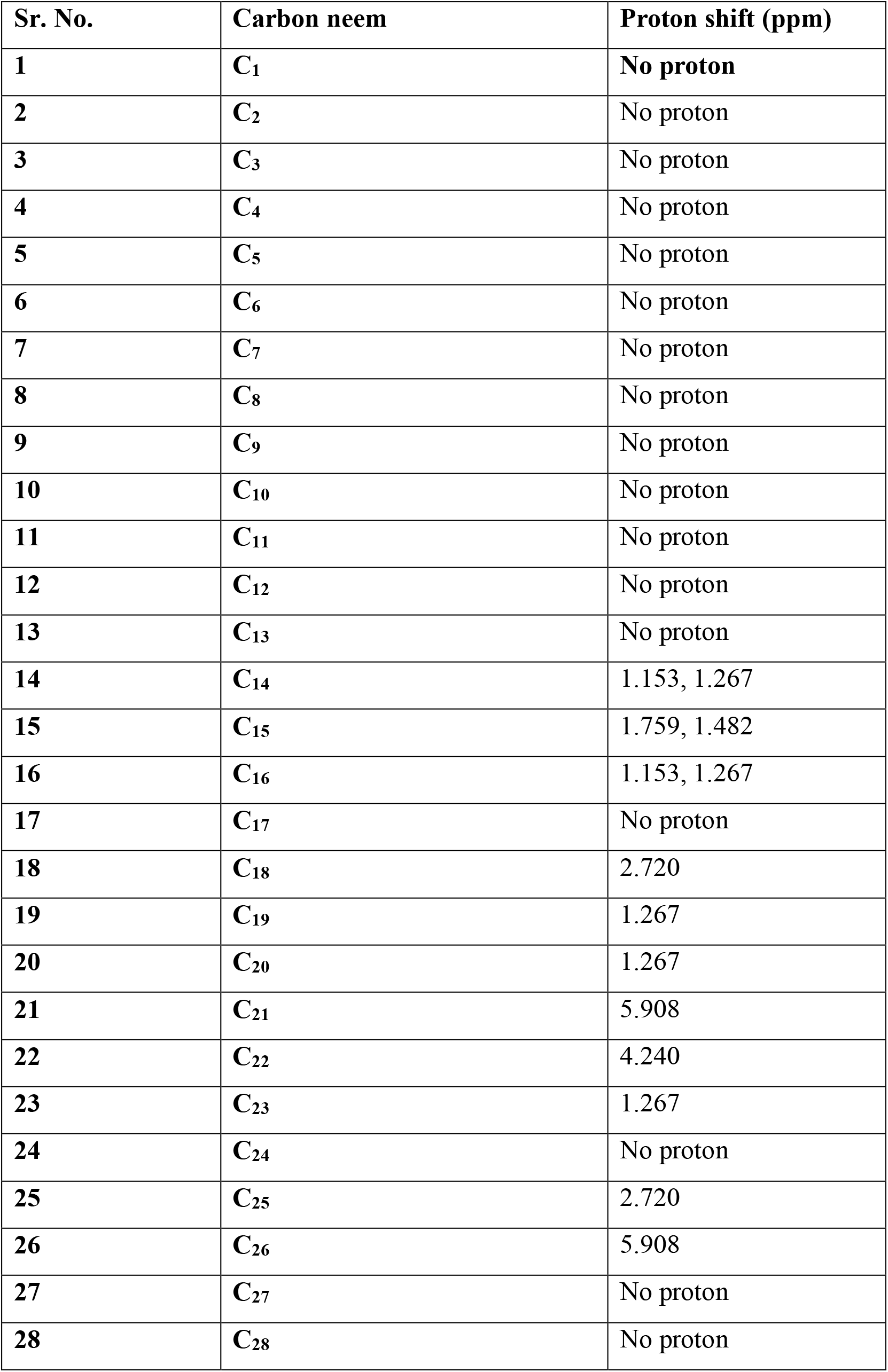
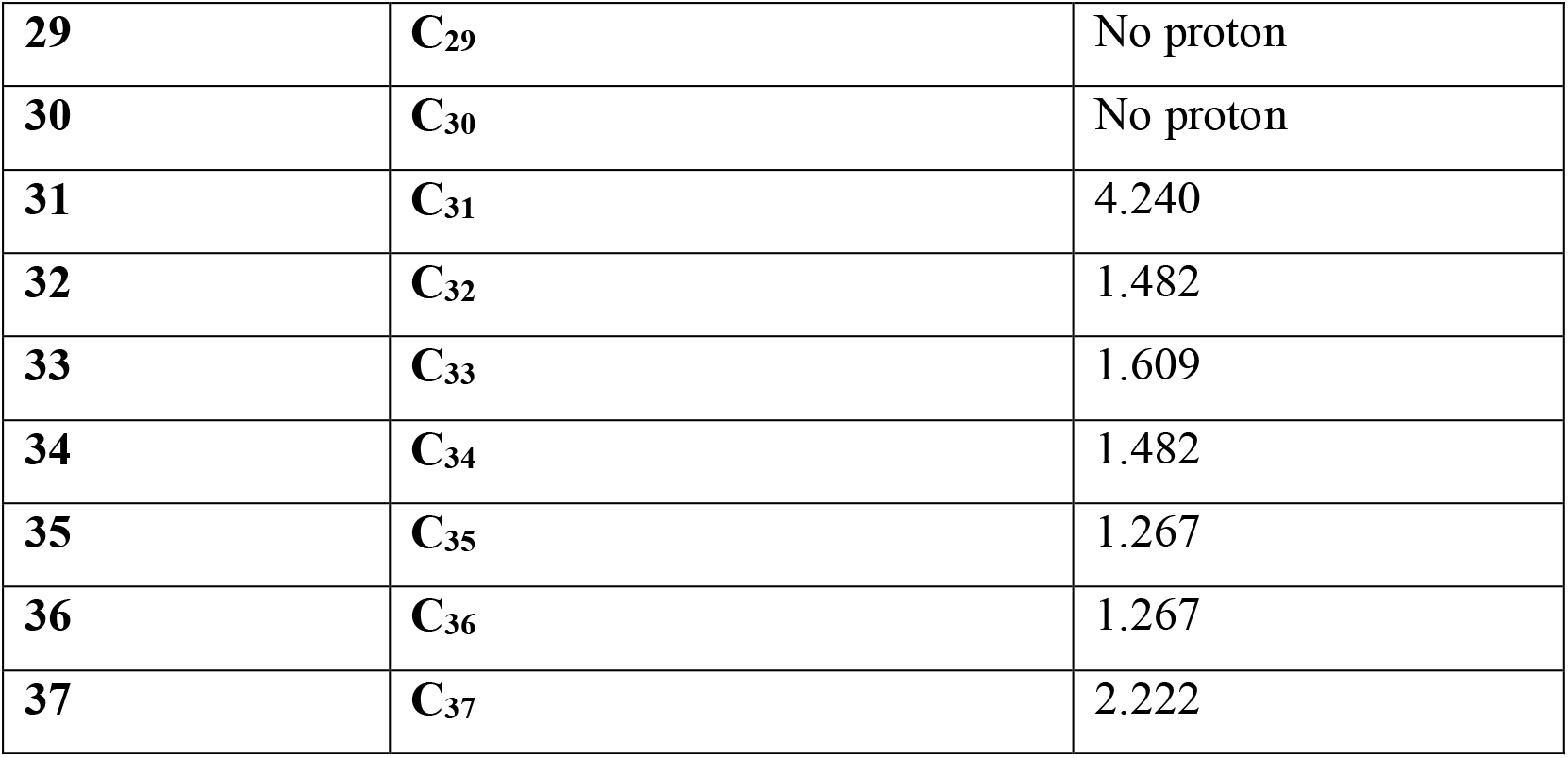
Proton shift position on carbon atom of *Azadirachta indica* fraction.

**Figure 4:**
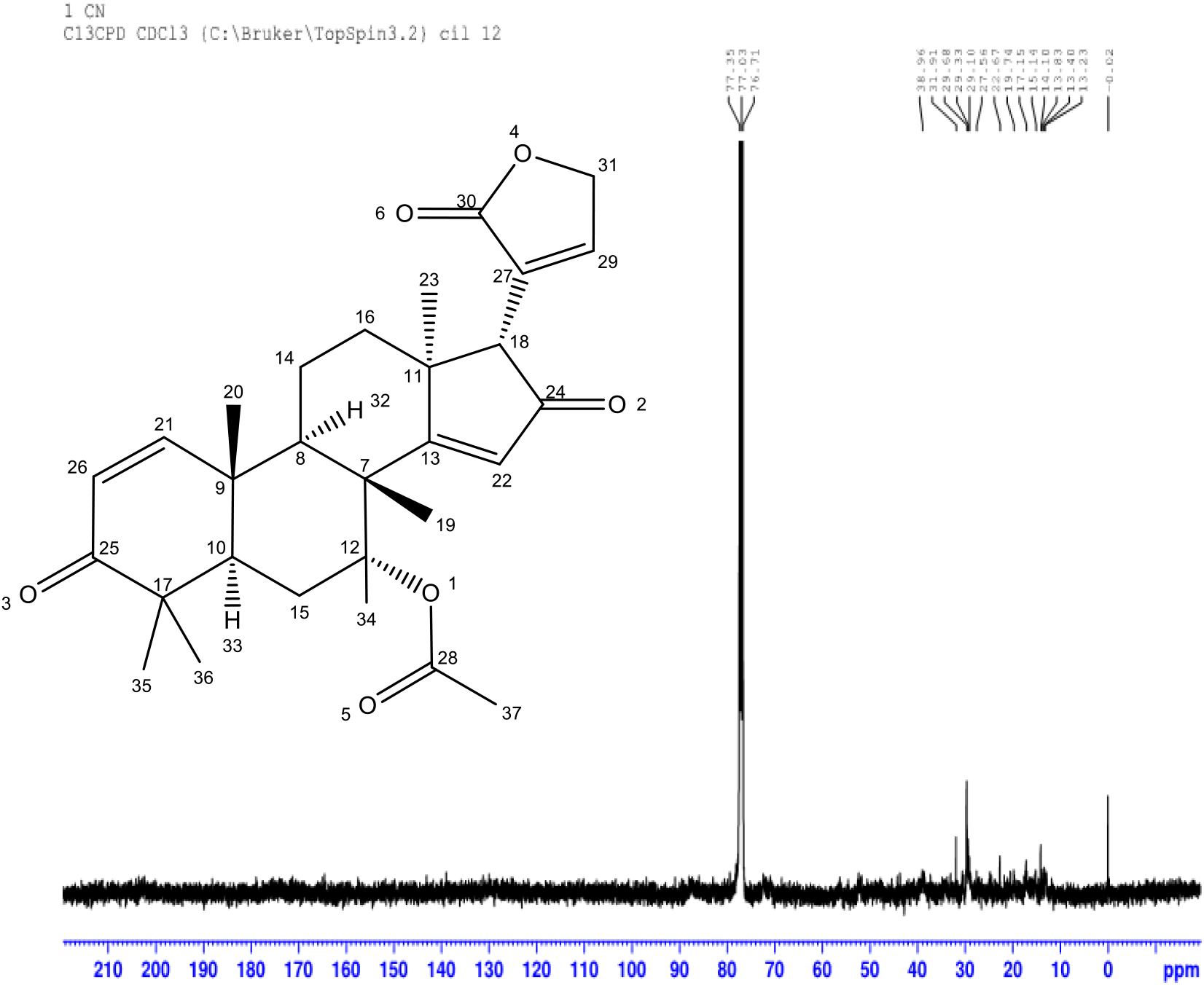
^C^NMR spectrum of different chromatographic peaks of *Azadirachta indica* fraction.

**Table 4:** Carbon shift position on carbon atom of *Azadirachta indica* fraction.

| Sr. No. | Carbon | Carbon shift (ppm) |
| --- | --- | --- |
| **1** | C<sub>1</sub> | No carbon |
| **2** | C<sub>2</sub> | No carbon |
| **3** | C<sub>3</sub> | No carbon |
| **4** | C<sub>4</sub> | No carbon |
| **5** | C<sub>5</sub> | No carbon |
| **6** | C<sub>6</sub> | No carbon |
| **7** | C<sub>7</sub> | 38.96 |
| **8** | C<sub>8</sub> | 38.96 |
| **9** | C<sub>9</sub> | 38.96 |
| **10** | C<sub>10</sub> | 38.96 |
| **11** | C<sub>11</sub> | 38.96 |
| **12** | C<sub>12</sub> | 76.71 |
| **13** | C<sub>13</sub> | 77.53 |
| **14** | C<sub>14</sub> | 15.14 |
| **15** | C<sub>15</sub> | 29.10 |
| **16** | C<sub>16</sub> | 29.68 |
| **17** | C<sub>17</sub> | 38.96 |
| **18** | C<sub>18</sub> | 38.96 |
| **19** | C<sub>19</sub> | 15.14 |
| **20** | C<sub>20</sub> | 19.74 |
| **21** | C<sub>21</sub> | 77.53 |
| **22** | C<sub>22</sub> | 77.53 |
| **23** | C<sub>23</sub> | 22.67 |
| **24** | C<sub>24</sub> | 77.53 |
| **25** | C<sub>25</sub> | 77.53 |
| **26** | C<sub>26</sub> | 77.53 |
| **27** | C<sub>27</sub> | 77.53 |
| **28** | C<sub>28</sub> | 77.53 |
| **29** | C<sub>29</sub> | 77.53 |
| **30** | C<sub>30</sub> | 77.53 |
| **31** | C<sub>31</sub> | 76.71 |
| **32** | C<sub>32</sub> | No carbon |
| **33** | C<sub>33</sub> | No carbon |
| **34** | C<sub>34</sub> | 19.74 |
| **35** | C<sub>35</sub> | 22.67 |
| **36** | C<sub>36</sub> | 22.67 |
| **37** | C<sub>37</sub> | 22.67 |

**Figure 5:**
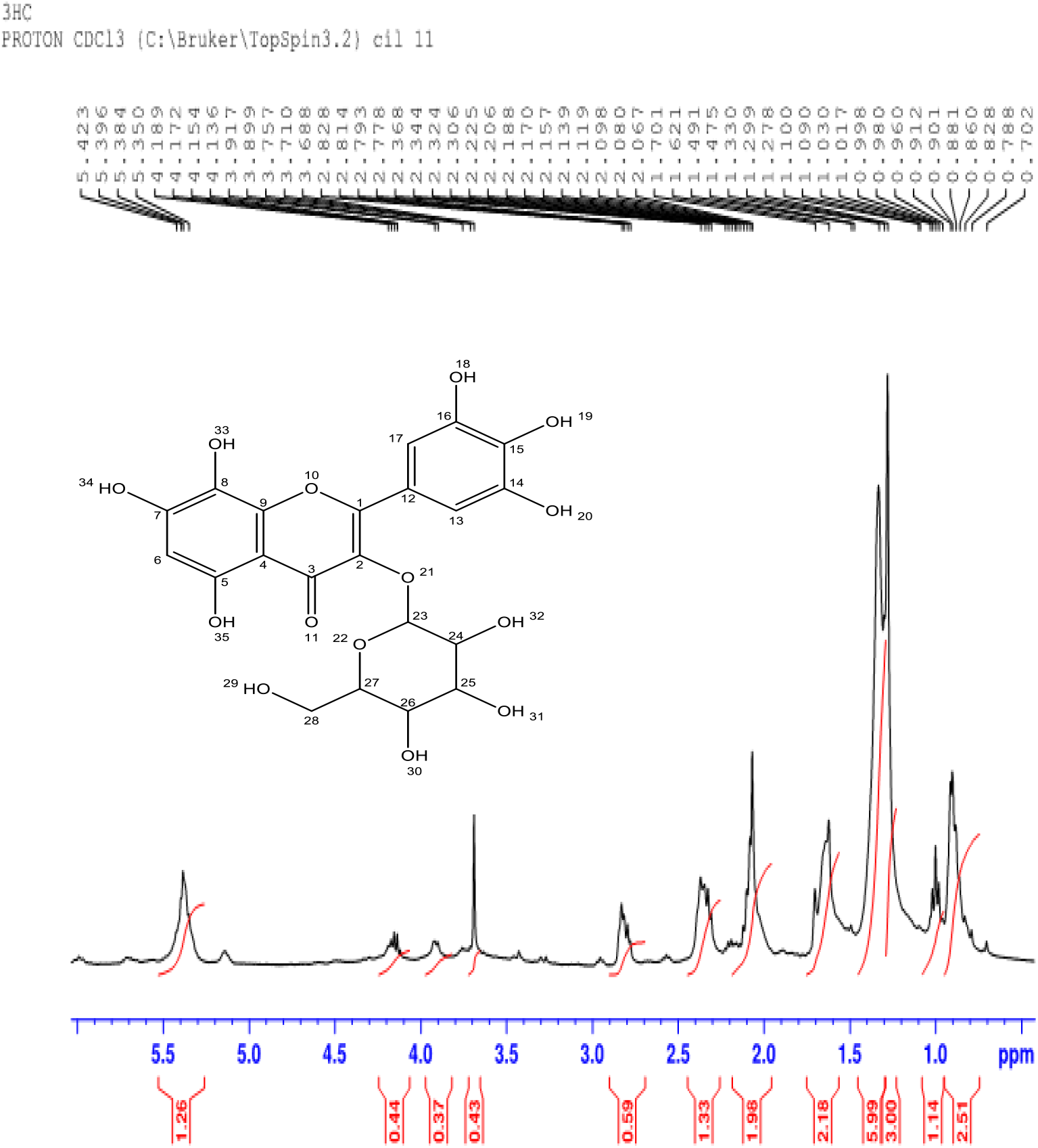
^H^NMR spectrum of different chromatographic peak of *Hibiscus sabdariffa* fraction.

**Table 5:**
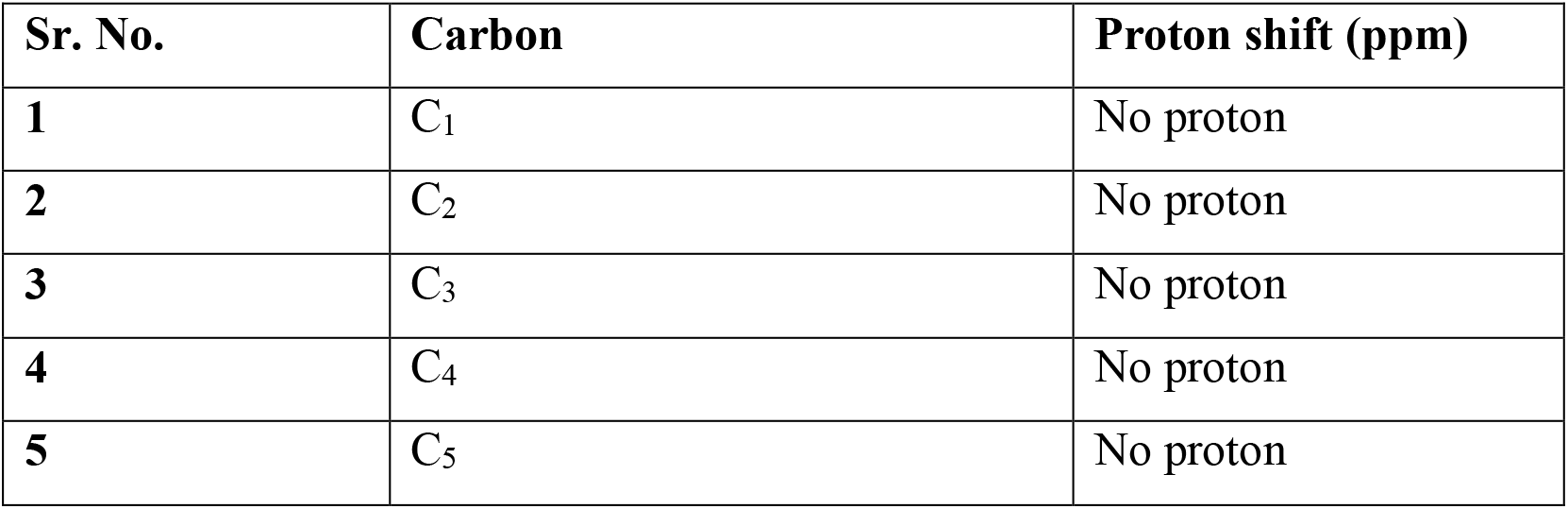
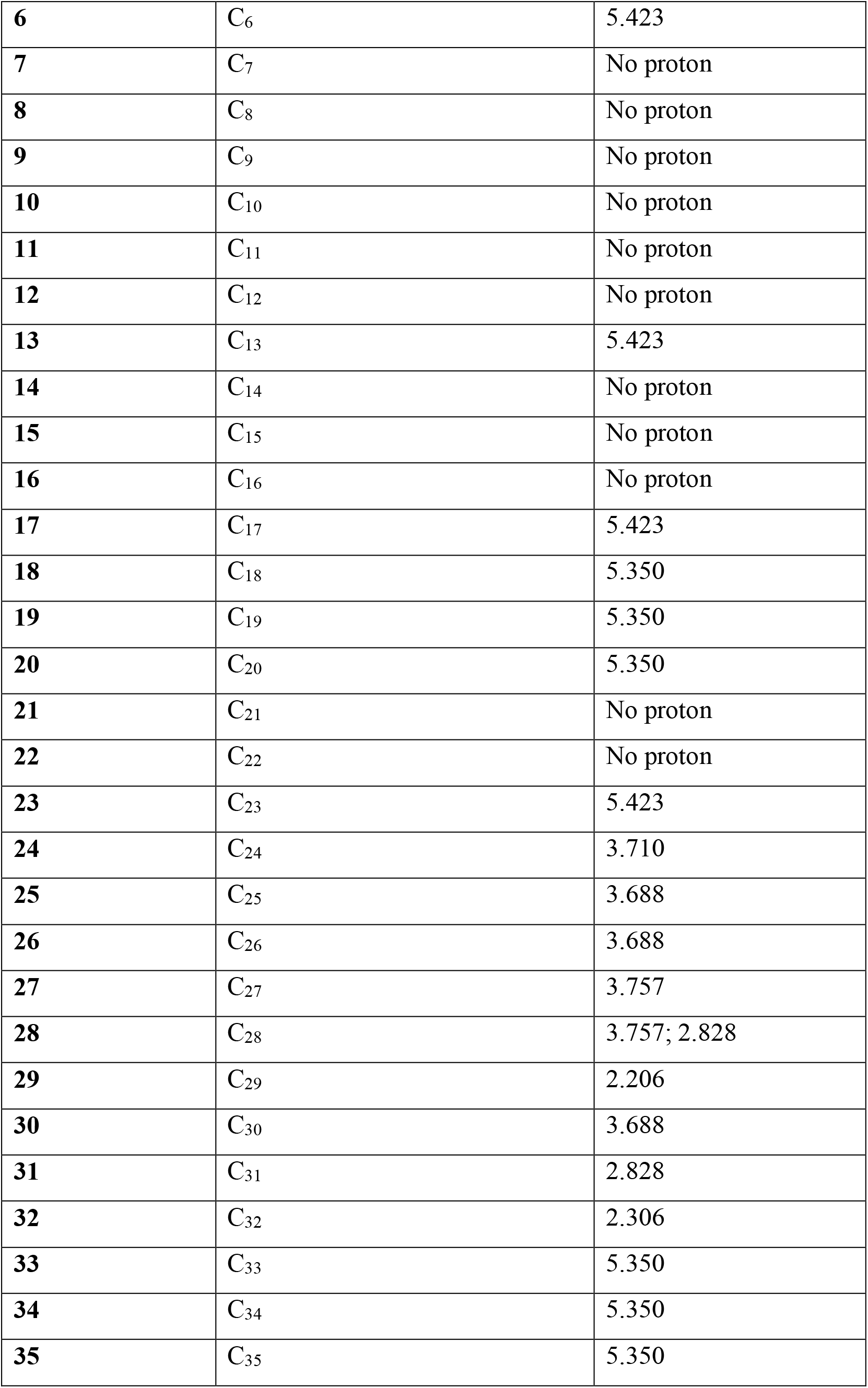
Proton shift position on carbon atom of *Hibiscus sabdariffa* fraction.

**Figure 6:**
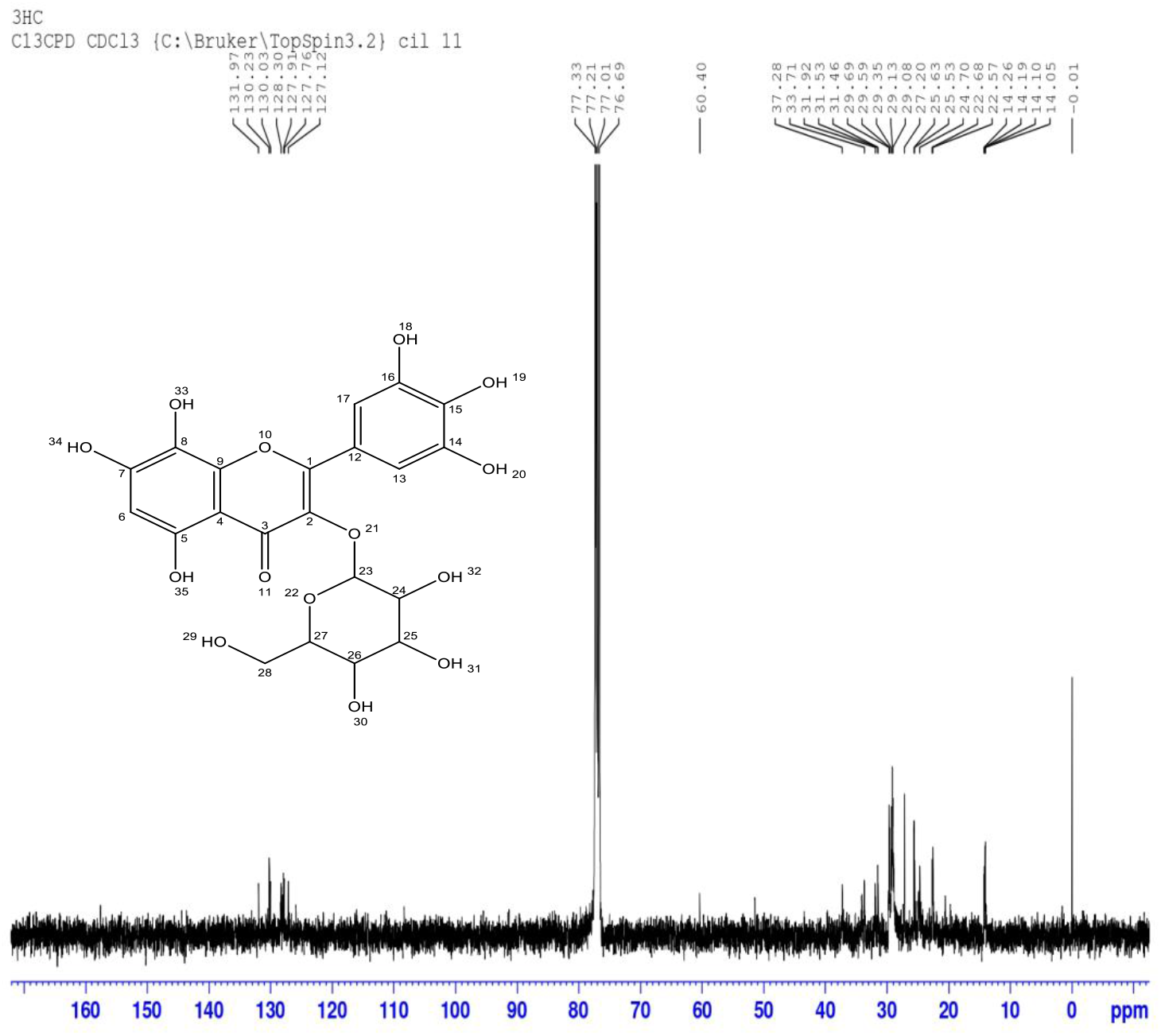
^C^NMR spectrum of different chromatographic peaks of *Hibiscus sabdariffa* fraction.

**Table 6:**
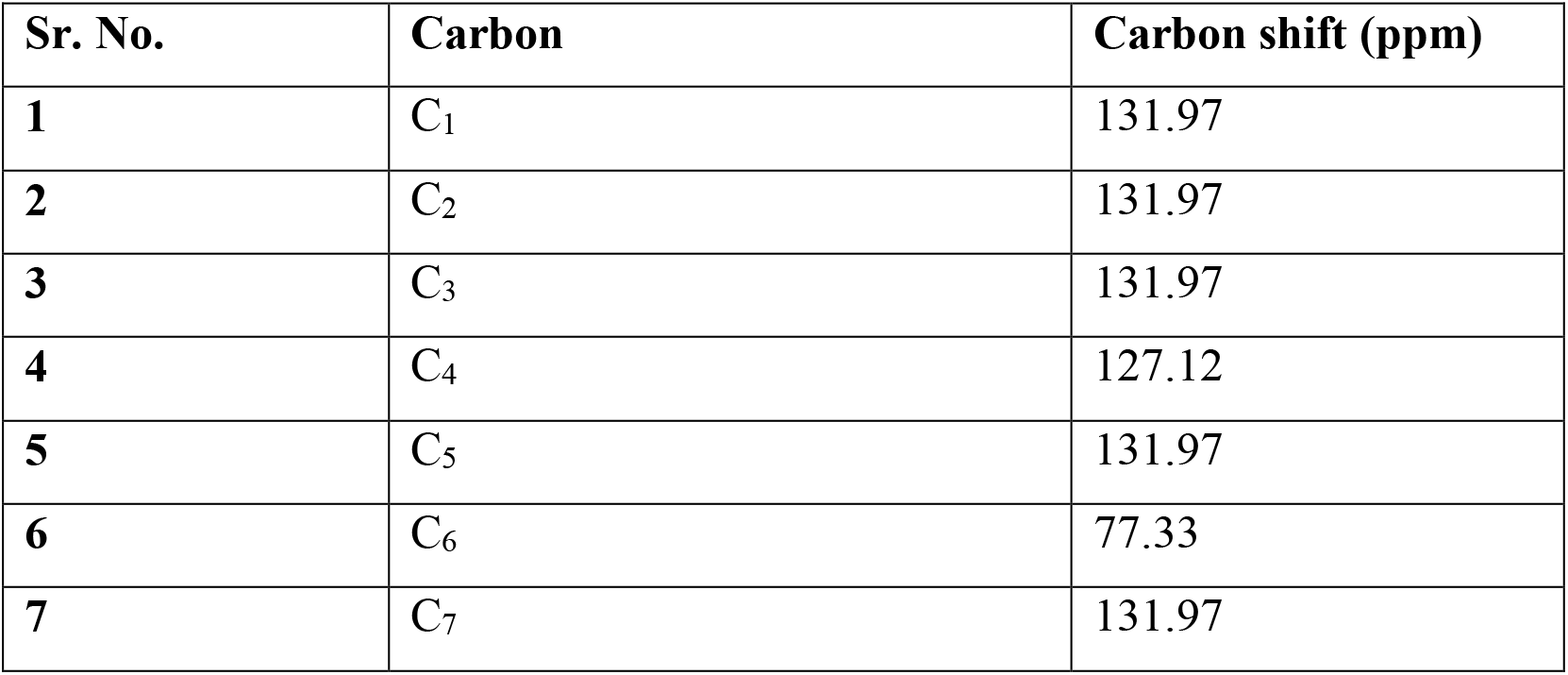
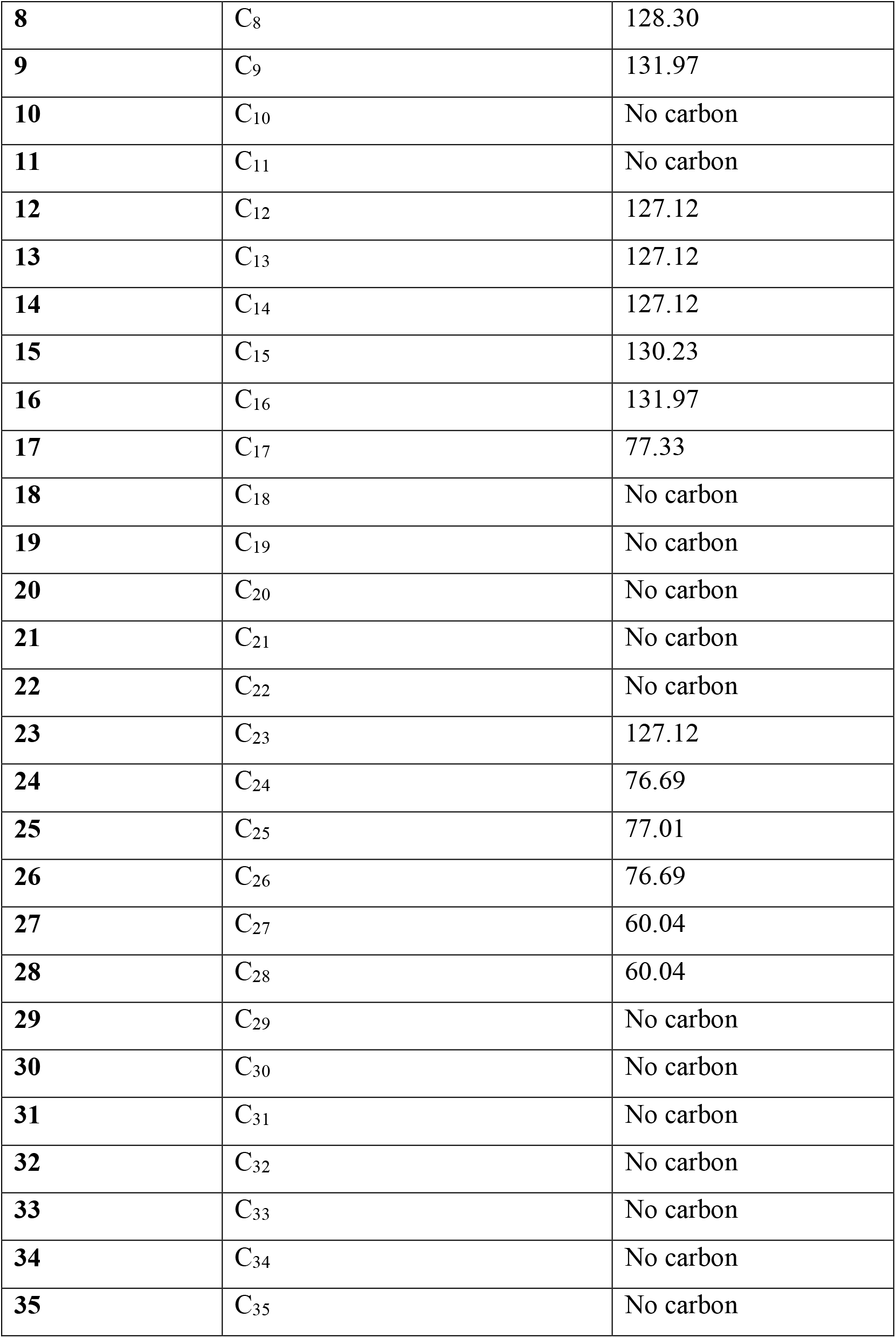
Carbon shift position on carbon atom of *Hibiscus sabdariffa* fraction.

